# *Akr1b7* functions as a master regulator in ovarian aging

**DOI:** 10.1101/2023.08.09.552567

**Authors:** Keishiro Isayama, Kenji Watanabe, Masato Ohtsuka, Seisuke Kimura, Tomoaki Murata, Takeshi Honda, Masataka Asagiri, Shun Sato, Hiroshi Tamura, Norihiro Sugino, Yoichi Mizukami

## Abstract

At the beginning of ovarian aging, the ovulation of immature oocytes is accelerated, leading to the arrest of ovulation despite the remaining oocytes. Here, RNA expression in the ovarian aging of mice is comprehensively analyzed during the estrous cycle after ovulation stimulation. The aldo-keto reductase *Akr1b7* pathway transiently activated in the ovaries of young mice disappears in those of old mice. *Akr1b7^—/—^* mice attenuate oocyte Akt activation essential for the follicular development in primordial follicles, and enhanced ovulation in immature oocytes. The estrous cycle is extended because of the prolonged diestrous stage by a sustained progesterone level in *Akr1b7^—/—^*mice ovaries, which is caused by the decline of *Cyp17a1*, a major metabolic enzyme of progesterone in *Akr1b7*-expressed theca cell layers. In summary, the decreased *Akr1b7* pathway causes ovulation of immature oocytes and a prolonged estrous cycle, typical symptoms of ovarian aging.

## Introduction

The ovary is a female-specific reproductive organ that critically functions as an oocyte supply.^1^ The oocyte is released from the mature follicle that forms a complex with the somatic cells according to the development of primordial follicles within the ovary.^2,3^ During follicular development, primordial follicles surrounding the monolayer of flat granulosa cells in the ovary develop into primary follicles with the proliferation of granulosa cells in response to follicle-stimulating hormone (FSH) and luteinizing hormone (LH) released from the pituitary gland.^1^ During the development of primary to secondary follicles, steroidogenic theca cells are recruited by oocyte-derived factors into the outer granulosa cells, forming multiple layers. Finally, the development of antral follicles induces the release of oocytes from mature follicles containing a fluid-filled cavity adjacent to the oocyte. After oocyte release, the follicles are transformed into corpora lutea (CLs), which secrete sex steroid hormones, estrogen and progesterone.^4^ The progesterone level is maintained by fertilization but rapidly decreases with regression of CLs in cases without fertilization.^5^ Primordial follicles are generated before birth, not during their lifetime. Notably, 1–2 million oocytes remain in the ovary at birth, decreasing to 300,000–400,000 primordial follicles at puberty and gradually declining during the reproductive age.^6,7^ Despite the cessation of ovulation, several hundred oocytes remain in the ovary during menopause, indicating that the cells surrounding the oocyte may play a critical role in ovarian aging. In clinical research on in vitro fertilization treatment, ovarian aging is known to cause poor ovulatory and steroidogenic responses to human chorionic gonadotropin (hCG) administration.^8,9^ Mice and rats have been proven to be models of ovarian aging in humans.^3^ Signs of ovarian aging, for example, a decreased number of developing follicles and a prolonged estrous cycle, were observed in mice aged approximately 28–48 weeks. Subsequently, cessation of the estrous cycle was observed at 44–72 weeks.^3,10^ Aged mice ovulated a few eggs and formed CLs in response to hCG stimulation, indicating that oocytes remained in the ovaries of aged mice. Because the inhibition of apoptosis in Bax-deficient mice contributes to the prolongation of ovarian life span, maintaining the number of follicles to inhibit ovarian aging might be a critical factor. The number of follicles is closely related to the menstrual cycle and is regulated by progesterone, FSH, and LH levels. Whole transcriptome analysis of single ovarian cells has been performed to elucidate the roles of oocytes and surrounding cells.^11-13^ The oxidative stress in the oocytes and granulosa cells was closely related to ovarian aging, impaired growth, and redox homeostasis.^13^ However, the signal transduction pathways and activated molecules that affect ovarian aging during folliculogenesis remain unclear in whole ovaries, where cell-to-cell interactions play essential roles. In this study, to elucidate the signaling pathway that affects ovarian aging, we comprehensively analyzed mRNA expression in mouse ovaries according to the time course during folliculogenesis. Sustained suppression of *aldo-keto reductase (Akr)1b7* was found in the ovaries of aged mice, in contrast with the transient expression in young mice. Using *Akr1b7^−/−^* mice deleted the start codon with an improved genome-editing via oviductal nucleic acid delivery (i-GONAD) method, we showed that AKR1B7 participated in the typical symptoms of ovarian aging, ovulation of immature oocytes, and a prolonged estrous cycle.

## Results

### Ovulation and follicular development in aged ovaries

The number of fetuses in female mice and a senescence marker were used to determine the age in weeks of mice with ovarian aging. Approximately 8 fetuses were in the 6-to 8-week-old female mice. Fetuses were rarely observed in mice older than 48 weeks. The estrous cycle was longer in 48-week-old mice than in 6-week-old mice, and the ovulated oocytes in 48-week-old mice included many immature oocytes (data not shown). OHdG, a marker of senescent cells, was detected in the cells surrounding the follicles of 48-to 56-week-old mice; however, staining was not observed in 6-week-old mice (Figure 1A).

**Figure 1.**
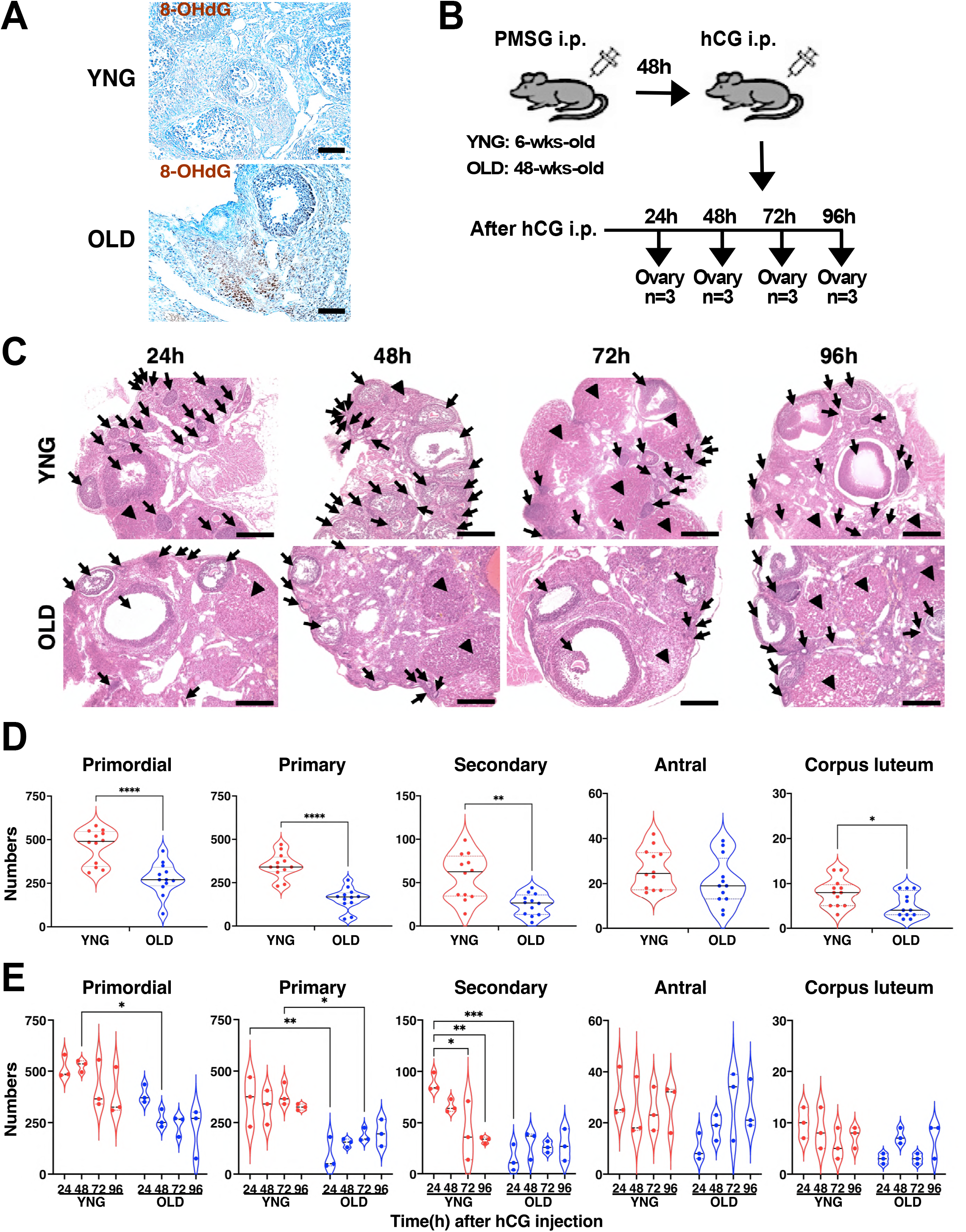
Follicular development in ovarian aging after ovulation stimulation. (A) Images show representative 8-OHdG stainings of sections in ovaries collected from YNG and OLD female mice. n = 3. (B) Illustrate shows sampling design of ovaries after ovulation stimulation. YNG and OLD female mice were injected intraperitoneally with 5IU-PMSG and 5IU-hCG; next, ovaries were collected at the indicated time. n = 3. Of the 2 ovaries, one was used for histological analysis, and the other was subjected to RNA expression analyses. (C) Representative H&E stainings of sections are shown in ovaries of YNG and OLD mice after PMSG/hCG injection. Arrows and arrowheads show corpora lutea and antral follicles, respectively. Scale bars, 500 µm. (D) Violin plots show total numbers of each follicle and corpora luteum using serial sections in YNG and OLD. n = 3. *P < 0.05, **P < 0.01, ****P < 0.0001; two-tailed Student’s t test. (E) Violin plots show numbers of each follicle stage and corpora luteum in serial sections in YNG and OLD after the PMSG/hCG injection. n = 3, *P < 0.05, **P < 0.01, ***P < 0.001; two-way ANOVA followed by Tukey’s multiple comparisons test.

Consistent with our data, a report revealed that the number of follicles and CLs was significantly less at 48 weeks of age than those at 8 weeks, although the follicles remained in the ovaries of 48-week-old mice.^14^ Based on these observations, 48-week-old mice were used as a model of ovarian aging, and the process of follicular development was examined during the estrous cycle after pregnant mare serum gonadotropin (PMSG) and hCG treatment in young (6-to 8-week-old) and old mice (48-to 56-week-old) (Figure 1B). There were more follicles in the ovaries of the young mice than in those of the old mice, except for antral follicles (Figures 1C and 1D). According to the time after hCG treatment, the number of each stage of follicles and CLs was slightly decreased in the ovaries of young mice (Figure 1E); the primordial follicles of the ovaries of the old mice continued to decline at 48 h after stimulation, but the number of follicles after the primary follicles more tended to increase than those at 24 h after stimulation (Figure 1E). Thus, follicular developmental dysfunction in aged ovaries may occur during the primordial follicle developmental stage.

### Whole transcriptome analysis during estrous cycle in ovarian aging

To elucidate the signaling pathways involved in follicular development dysfunction in aged mice, whole transcriptome analysis was performed on mouse ovaries during the estrous cycle after ovulation treatment. More than 15,000 genes were detected in approximately 30 million reads in each ovary, and the expression of the representative genes almost corresponded to that obtained by quantitative polymerase chain reaction (PCR; Figure S1). Principal component analysis (PCA) revealed that gene expression patterns in the ovaries of young and old mice were clearly separated by the first principal component (PC1), independent of the estrous cycle. Gene expression in the ovaries of young mice, according to the time course after ovulation stimulation, shifted in a positive direction on the second principal component (PC2), and the ovaries of old mice at 96 h returned to a negative direction on PC2 (Figure 2A). The marker genes in the steps of follicular development using the single-cell dataset^13,15,16^ demonstrated that the genes (*Ere*g, *Adamts*1, and *Edn2*) involved in ovulation in young mice showed transiently high expression 24 h after stimulation, and those in old mice were almost constant during the cycle (Figure 2B and 2C). The marker genes for primordial (*Foxo3*, *Fos*, and *Ddx4*) and primary follicles (*Figla*, *Bmp15*, and *Gdf9*) and theca cells (*Cyp17a1*, *Col1a2*, and *Insl3*) continued to be expressed at high levels in the ovaries of young mice, but during the estrous cycle in old mice, sustained expression at low levels was observed (Figure 2B and 2C). CG-responsive genes *Cyp11b1* and *Ereg* were rapidly decreased 48 h after stimulation, and *Tgfb1* was immediately induced by stimulation (Figure 2D). The responsive genes in old mice were slightly regulated or were not responsive to CG (Figure 2B). The alteration of gene expression during the estrous cycle was observed in young mice at 48 hr after stimulation, and the alternation was obviously weak in old mice (Figure 2D and 2E). The 141 transiently upregulated genes at 24 h after stimulation were analyzed using ingenuity pathway analysis (IPA), and CG was detected as the first candidate upstream factor (Figure 2F). The downstream effectors indicated that steroid hormone pathways played a central role in follicular development in response to CG (Figure 2G). Network analysis illustrated that ovulation stimulation with CG and LH was closely associated with Aldo-keto reductases AKR1B7 and AKR1B8 (Figure 2H). The expression of *Akr1b7* transiently increased 24 h after ovulation stimulation, and its expression rapidly decreased at 48 h, although *Akr1b8* was almost constant during the estrous cycle (Figure S2). CG and FSH were detected again in the network analysis at 96 h after stimulation, indicating that the estrous cycle entered the next cycle (Figure S3A–C). In old mice, the CG-activated NADPH oxidase pathway was detected in the downregulated genes, and CG and FSH were not detected until 96 h, except after exogenous administration, in contrast with the pathway detected in young mice (Figure S3D–F). These observations indicate that decreased *Akr1b7* expression may be implicated in follicular developmental dysfunction during ovarian aging.

**Figure 2.**
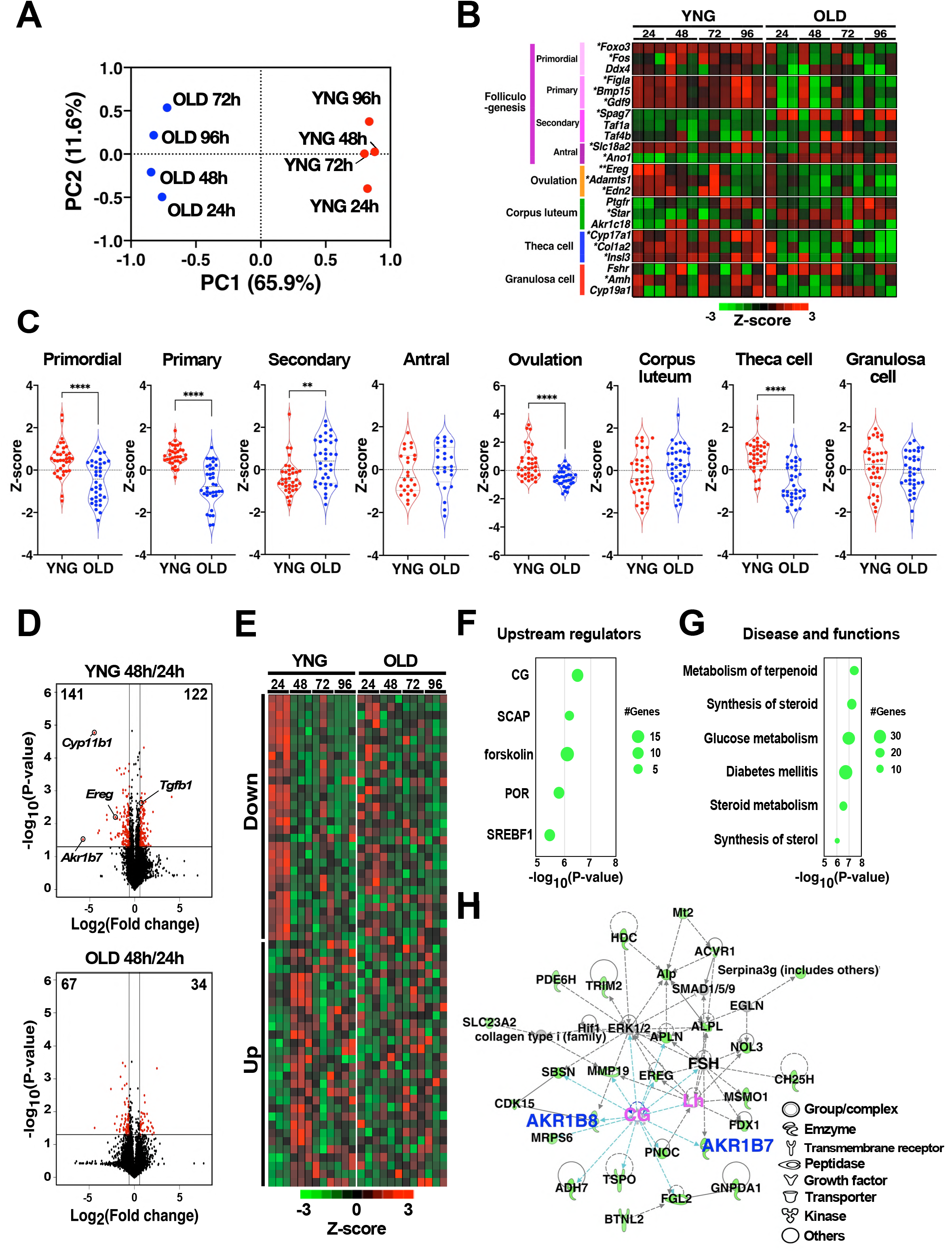
Whole transcriptome analysis in ovarian aging after ovulation stimulation. (A) Principal component analysis using mRNA expression values in ovaries of YNG and OLD (n = 3) was plotted using the values calculated from PC1 and PC2. (B) CPM in the indicated marker genes transformed to log_2_ and then normalized by z-score are shown on heatmap. (C) Z-scored values of marker gene expressions indicated in Figure 2B were plotted by each group. (D) Volcano plot shows differentially expressed genes in the ovaries from 24 to 48 h after PMSG/hCG injection. The negative log_10_-transformed P values are plotted against the log_2_ transformation of fold changes. The genes with a fold change of more than 1.5 and P value below 0.05 are plotted in red circles. (E) Heat maps show z-score of genes indicated in red circles of Figure 2D. (F) The gene set detected as downregulated genes in Figure 2D was analyzed by IPA pathway software, and the detected upstream regulators are shown. (G) The gene set detected as downregulated genes in Figure 2D was analyzed by IPA, and the pathways related to disease and functions are shown. (H) The gene set detected in Figure 2D was analyzed by IPA software, and the representative pathway is shown.

### *Akr1b7* mRNA expression in ovarian aging

The expression of *Akr1b7* mRNA during follicular development was confirmed using quantitative PCR. The expression was detected at a high level 24 h after hCG administration and rapidly decreased during the next 24 h, consistent with the data from the whole transcriptome analysis (Figure 3A). *Akr1b7* expression in aged mice was low after stimulation, and the low expression persisted during the estrous cycle. Immunohistochemical staining using an antibody against human AKR1B10, with an epitope sequence identical to that of mouse AKR1B7, was performed in both the theca cell layer and stromal cells after ovulation stimulation (Figure 3B). The staining in the theca cells expressing the LH and CG receptors decreased significantly 48 h after stimulation (Figure 3C), similar to mRNA expression. To confirm the staining detected in the theca cells, we examined the expression in the target regions extracted from ovarian tissue slices via laser microdissection using superfamily-specific primers. Among the *Akr1b* superfamily members, *Akr1b7* and *Akr1b8* mRNAs were exclusively expressed in the ovaries of young mice (Figure 3D and S2). *Akr1b7* was mainly expressed in the theca cell layer of the ovaries of young mice, consistent with the expression of *Vimentin (Vim)*, a marker of theca cell layers, and its expression contrasted with that of *anti-Müllerian hormone (Amh)*, a marker of granulosa cells. The expression of *Akr1b8* was observed in both granulosa and theca cell layers (Figure 3E). *Akr1b7* mRNA levels in the theca cell layers of old mice were significantly lower than those in young mice (Figure 3F). Staining with an anti-human AKR1B10 antibody corresponded with *Akr1b7* mRNA expression, indicating that the antibody recognizes mouse AKR1B7 in the ovary. Isocaproaldehyde (ICA) and 4-hydroxynonenal (4-HNE), known as substrates for AKR1B7, were used to examine the enzymatic characteristics of AKR1B7 in ovaries. The apparent Km values for ICA and 4-HNE were 550 µM and 36 µM, respectively, in the ovaries of young mice (Figure 3G), and the substrate affinities were almost the same as the Km values using recombinant AKR1B7 for ICA and 4-HNE at 320 µM and 62 µM, respectively.^17^ In the ovaries of old mice, the reduction activities of ICA and 4-HNE were slightly lower than those in young mice, and the reduction magnitude differed from that of *Akr1b7* mRNA in old mice (Figure 3H). AKR1B8 may compensate for the detoxification activity in the ovaries because AKR1B8 has enzymatic activities for ICA and 4-HNE.^18^ Prostaglandin (PG)F_2α_-forming activity was measured in the ovaries after hCG treatment because AKR1B7 has the transforming activity of PGH_2_ to PGF_2α_, and AKR1B8 has no activity for the substrate. PGF_2α_-forming activity in the ovaries of old mice was significantly lower than that in the ovaries of young mice (Figure 3I), and PGF_2 α_ production was positively correlated to *Akr1b7* mRNA levels with a correlation coefficient of 0.73 (Figure 3J). These observations indicate that the decrease in *Akr1b7* mRNA during ovarian aging was at least partially associated with a reduction in PGF_2α_-forming activity.

**Figure 3.**
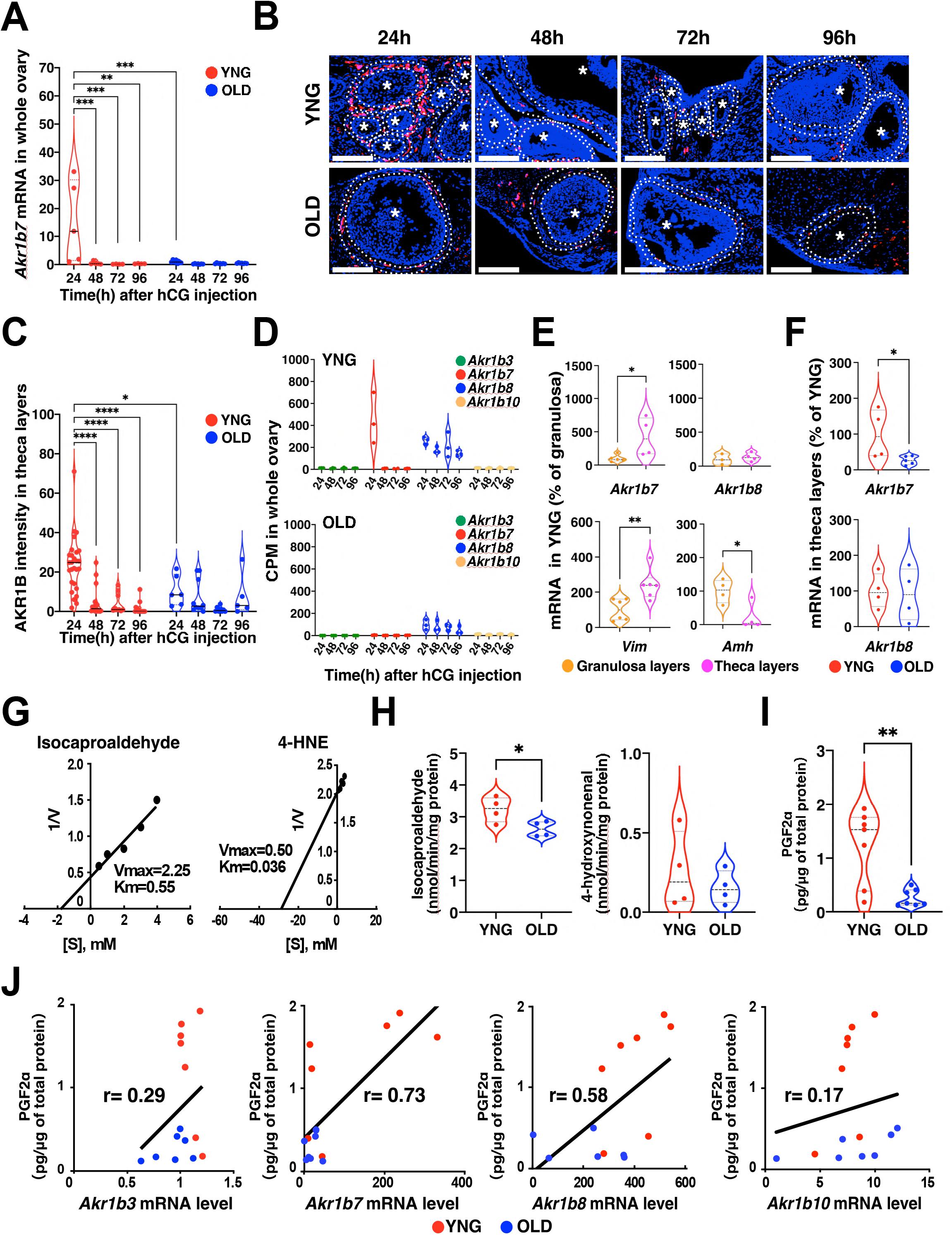
Expression and enzymatic analysis of AKR1B7 in ovarian aging. (A) Quantitative PCR analysis of *Akr1b7* mRNA expression was conducted using ovaries collected from YNG and OLD female mice at the indicated time after PMSG/hCG injection. Violin plots show the expression level relative to *Gapdh*. n = 4–7, **P < 0.01, ***P < 0.001; two-way ANOVA followed by Tukey’s multiple comparisons test. (B) Images show immunohistochemical staining with anti-human AKR1B10 antibody having identical epitope sequence for AKR1B7 in the ovaries of YNG and OLD female mice at the indicated time after PMSG/hCG injection. The sections were incubated with the antibody and were detected with a DAB reaction. Images of stained section were pseudocolored with DAB stains (red), and with hematoxylin counterstains (blue). Circle with dotted lines represents the theca layer. n = 3, Scale bars (white line), 100 µm. (C) Mean intensities of the staining in theca cell layer in Figure 3B were calculated by dividing the signal intensity by the area. Number of follicles was 5–26. n = 3, *P < 0.05, ****P < 0.0001; 2-way ANOVA followed by Tukey’s multiple comparisons test. (D) Violin plots show expressions of AKR1B superfamily genes determined using whole transcriptome analysis. n = 3. (E) Violin plots show *Akr1b7* and *Akr1b8* mRNA relative expression to *Actb* in laser micro-dissected theca cell and granulosa cell layer of YNG and OLD female mice. *Vim* and *Amh* mRNA were measured as a marker gene of theca and granulosa, respectively. n = 3–6, *P < 0.05, **P < 0.01; two-tailed Student’s t test. (F) Violin plots show *Akr1b7* and *Akr1b8* mRNA relative expressions to *Actb* in theca layers of YNG and OLD. *P < 0.05; two-tailed Student’s t test. (G) Lineweaver-Burk plot was determined from the reductase activities for ICA (left panel) or 4-HNE (right panel) in YNG ovary at 24 h after PMSG/hCG injection. (H) Violin plots show reductase activities for ICA (left panel) or 4-HNE (right panel) in the cytosol extracts of YNG and OLD ovaries at 24 h after PMSG/hCG injection. n = 4, *P < 0.05; two-tailed Student’s t test. (I) PGF_2α_ concentrations were quantified by ELISA using the ovary homogenates of YNG and OLD at 24 h after PMSG/hCG injection. n = 7, **P < 0.01; two-tailed Student’s t test. (J) mRNA expression levels of the indicated *Akr1b* superfamily were plotted against the PGF_2α_ concentration in YNG (red) and OLD (blue). n = 7; r indicates Pearson correlation coefficient.

### Generation of *Akr1b7*-deficient mice with i-GONAD

We produced *Akr1b7* locus-disrupted mice (*Akr1b7^−/−^*) by using genome editing for preimplantation embryos within the oviduct with electroporation, namely, i-GONAD to elucidate the physiological roles in AKR1B7 in ovarian aging (Figure 4A). Guide RNA (gRNA) was designed for the region, including the protospacer adjacent motif sequence 5 bases from the *Akr1b7* start codon, and was inserted into the oviduct with the CAS9 protein (Figure 4B). Fifty-two fetuses were born from 6 female mice treated with i-GONAD, and both alleles of *Akr1b7* were disrupted around the start codon in 13.5% of the fetuses. Twenty fetuses (23.1%) had a hetero-deletion mutation within the *Akr1b7* locus, and 13 fetuses (25.0%) exhibited mosaicism (Figure 4C). The representative genome sequence of mice with disrupted *Akr1b7* had 20 bases deleted, including the start codon of *Akr1b7* (Figure 4D). In the ovaries of young wild-type (WT) mice, 24 h after hCG stimulation, the reads of *Akr1b7* in the whole transcriptome analysis were mapped uniformly to the whole exon of *Akr1b7*. The mapped reads in *Akr1b7^−/−^*mice were not detected around the start codon of *Akr1b7* exon 1 (Figure 4E), although the reads were detected in all regions except for the start codon. The AKR1B7 staining mainly detected in the theca cell layer of WT mice was unobserved in *Akr1b7^−/−^* mice (Figure 4F and 4G). The *Akr1b7^−/−^*mice had no effects on the expressions of the other *Akr1b* superfamily (Figure S4). These observations indicate that *Akr1b7*-disrupted mice were successfully generated using the i-GONAD method.

**Figure 4.**
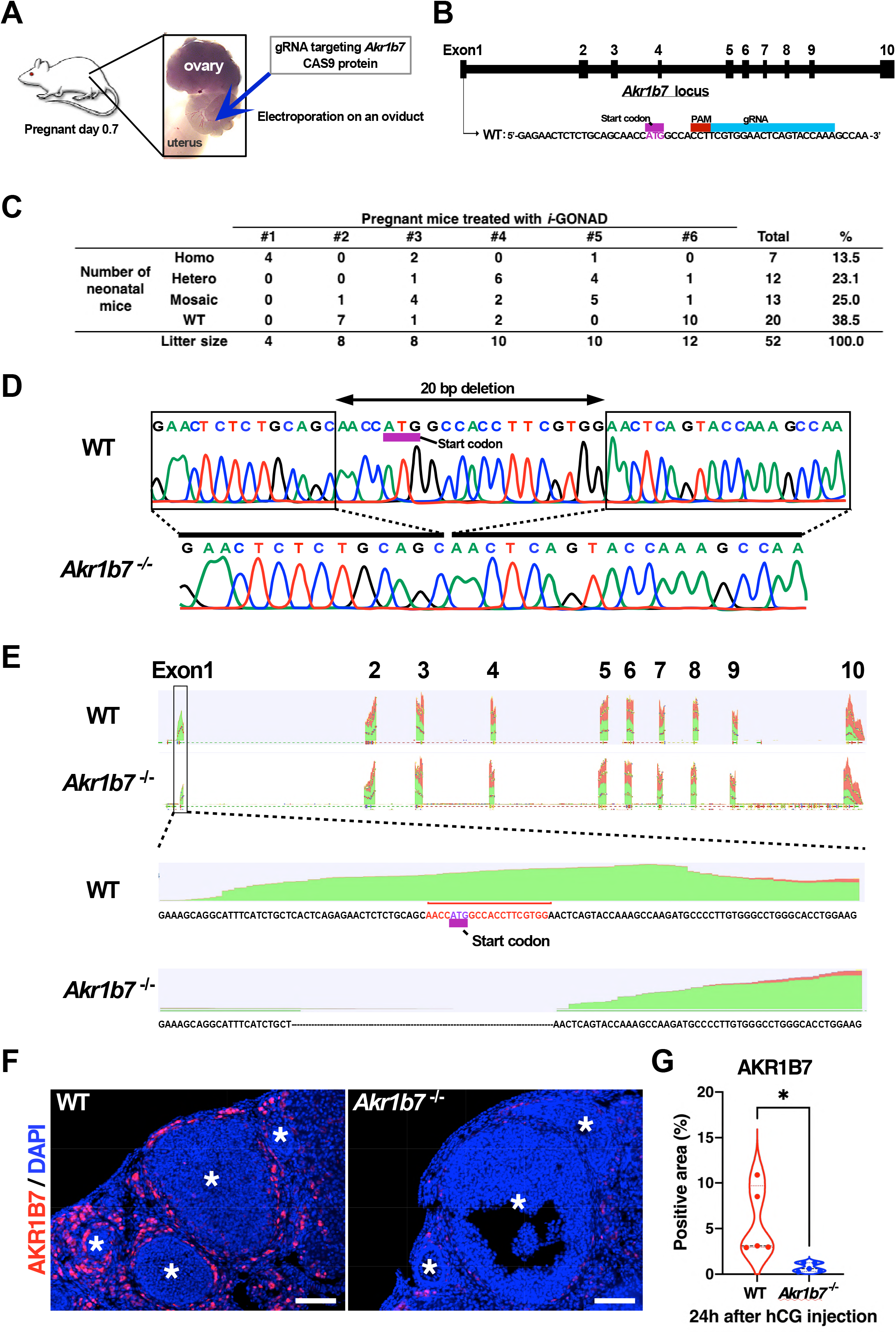
Generation of *Akr1b7*-deficient mice with i-GONAD method. (A) The gRNA targeting around the start codon of *Akr1b7* and CAS9 protein was injected and electroporated on the oviduct of ICR female mice at pregnant day 0.7. (B) Genomic *Akr1b7* locus edited by the complex of *Akr1b7*-gRNA and CAS9 is shown. (C) Table shows the genotyping in pregnant mice treated with i-GONAD method. (D) Chromatograms by Sanger sequencing show DNA sequences of PCR products amplified from genomic DNA, including the *Akr1b7* gRNA target site. Peaks on the chromatograph, green for A, red for T, blue for C, and black for G. (E) Mapping data show reads mapped on whole exon regions of *Akr1b7* with RNA-seq in WT and *Akr1b7^−/−^*(upper panel), and the data are enlarged in exon1 region of *Akr1b7* (lower panel). (F) Sections of immunofluorescence show staining of AKR1B7 in ovaries collected from WT and *Akr1b7^−/−^* female mice at 24 h after PMSG/hCG injection. Asterisks indicate a growing follicle belonging to the secondary or antral stage. Scale bars, 200 µm. (G) Violin plot show AKR1B7-positive area (%) in total area of an ovarian section in Figure 3F. n = 5, *P < 0.05; two-tailed Student’s t test.

### Ovulation of immature oocytes in *Akr1b7^−/−^* mice

*Akr1b7^−/−^* YNG mice and WT YNG mice had no significant differences in the morphology of developmental follicles observed using H&E staining (Figure 5A). The litter size and number of ovulated oocytes using identical 6-week-old mice of WT, *Akr1b7^+/–^*, and *Akr1b7^−/−^*were similar in all groups (Figure 5B and 5C). To examine the involvement of *Akr1b7* in follicular development, we comprehensively analyzed mRNA expression in the ovaries of 6-week-old *Akr1b7^−/−^* mice. The marker genes in follicular development classified by scRNA-seq analysis illustrated that the expression of genes associated with ovulation was significantly increased in *Akr1b7^−/−^*mice (Figure 5D and 5E). Furthermore, the oocytes ovulated in *Akr1b7^−/−^* mice were smaller than those in WT mice and were immature oocytes in the germinal vesicle and in the metaphase I stage without the first polar body in the structures, similar to the aged mice (Figure 5F and 5G). Immature oocytes comprised approximately 70% of the ovulated oocytes (Figure 5H). The analysis of follicular stages in old mice showed that primordial follicles were significantly decreased in 16-week-old mice, accompanied by more CLs in *Akr1b7^−/−^* mice than in WT mice (Figure 5I).

**Figure 5.**
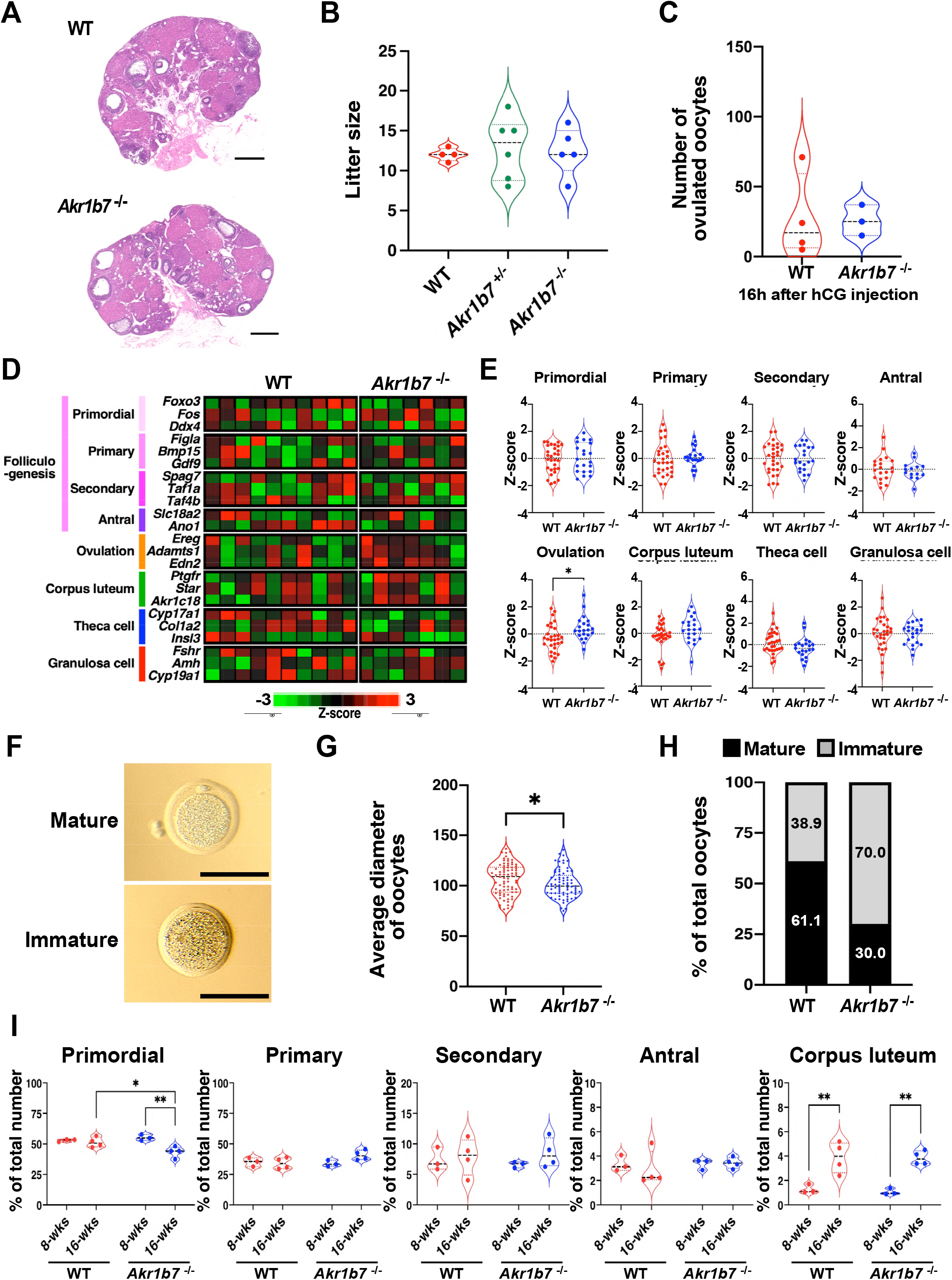
Follicular development in *Akr1b7*-deficient mice after ovulation stimulation. (A) Slices with H&E staining show the ovaries of YNG WT and *Akr1b7^−/−^* mice. (B) Violin plots show litter sizes of WT, *Akr1b7^+/–^*, *Akr1b7^−/−^* mated mice at 8-week-old. n = 4–6. (C) The number of ovulated oocytes obtained from YNG mice was counted at 16 h after PMSG/hCG injection. n = 3–4. (D) Heatmap shows z-scored values of the indicated marker gene expressions. WT; n = 10, *Akr1b7*^−/−^; n = 7. (E) Violon plots show z-scores of marker gene set in Figure 5D. (F) Images show representative images of oocytes obtained from WT (upper) and *Akr1b7^−/−^* mice (lower). Scale bars, 100 µm. (G) Violin plots show average diameters of an oocyte, including a zona pellucida. WT; n = 79, *Akr1b7*^−/−^; n = 72. (H) Bar graphs show rates of mature and immature oocytes in total oocytes. WT; n = 90, *Akr1b7*^−/−^; n = 70. (I) Violin plots show the rates of follicles at each stage in all follicles. Each follicle and corpus luteum were counted in serially ovarian sections in 8-week-old and 16-week-old mice of WT and *Akr1b7*^−/−^ at 24 h after PMSG/hCG injection. n = 3 in 8-week-old, n = 4 in 16-week-old. *P < 0.05, **P < 0.01; two-way ANOVA followed by Tukey’s multiple comparisons test.

### Downstream molecules of AKR1B7 in follicular development

To elucidate the signaling pathway of the folliculogenesis activated by AKR1B7, we performed PCA by using an RNA-seq analysis of the ovaries of *Akr1b7^−/−^* mice. PCA plots revealed a clear difference in gene expression between WT and *Akr1b7^−/−^* mice on PC7 (Figure 6A), and the loading factors that contributed to the separation of PC7 were used for IPA pathway analysis. The activated genes in *Akr1b7^−/−^* mice indicated that CG is an upstream molecule, suggesting that ovulation stimulation was enhanced in *Akr1b7^−/−^* mice (Figure 6B). The effector pathways in *Akr1b7^−/−^* mice showed that glucose metabolism disorders and diabetes mellitus were upregulated (Figure 6C). The analysis of the downregulated genes indicated the inhibition of glucose and lipid metabolism and a decrease in steroid-associated transcription factors (Figures 6D and 6E). The pathways detected in *Akr1b7^−/−^* mice were similar to those detected in the ovaries of old mice (Figures 2F and 2G). Ovarian glucose metabolism is closely associated with the PI3K/Akt signaling pathway, which is involved in folliculogenesis dysfunction, as observed in ovarian aging.^19^ Akt was significantly activated in the granulosa cell layers of developing follicles and in the oocytes of the primordial follicles of WT mice after hCG treatment, and Akt activation in these regions was attenuated in *Akr1b7^−/−^* mice (Figure 6F–6H, and Figure S5A–D). In primordial follicles, extracellular signaling molecules are necessary to activate Akt by *Akr1b7* restrictedly expressed in stromal cells, and the tissue distribution of the c-Kit ligand (KITL) that activates Akt was examined.^20^ KITL staining was observed in the cytoplasm of *Akr1b7*-expressed stromal cells, in addition to the punctate staining of granulosa cells, and only the KITL staining of *Akr1b7*-expressed cells disappeared in *Akr1b7*^−/−^ mice (Figure 6I). At 16 weeks of age, the KITL staining of granulosa cells was attenuated in *Akr1b7*^−/−^ mice, which was consistent with a decrease in primordial follicles (Figure S5E). *Akr1b7* regulated the Akt activation necessary for follicle development through KITL in stromal cells.

**Figure 6.**
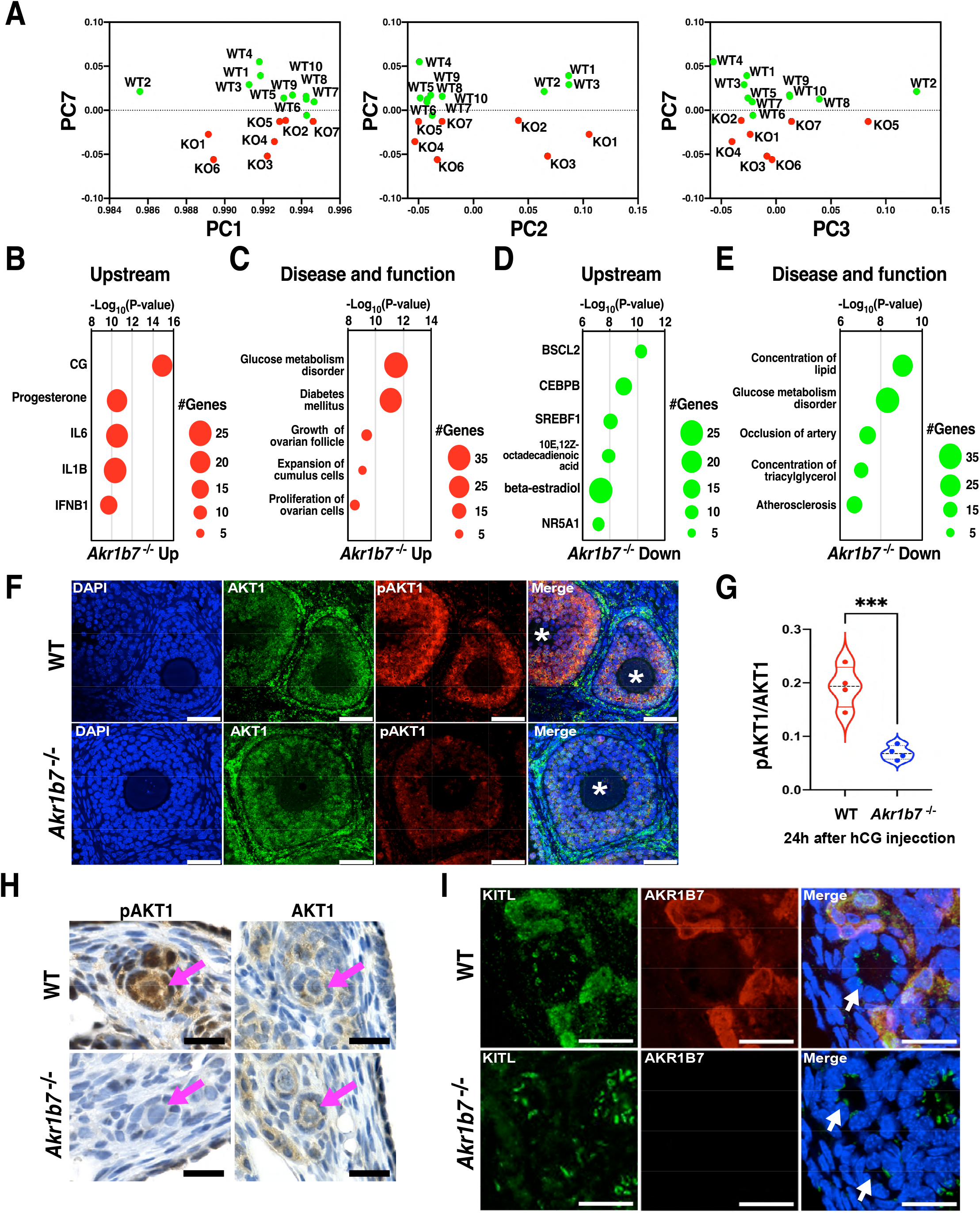
Signal transduction in primordial follicles of *Akr1b7*-deficient mice after ovulation stimulation. (A) Principal component analysis was carried out using mRNA expression of ovaries collected from YNG WT (n = 7) and *Akr1b7*^−/−^ mice (KO) (n = 10) at 24 h after PMSG/hCG injection. Dot plots show the values of PC1 (left panel), PC2 (center panel), or PC3 (right panel) against the values of PC7. (B–E) The upregulated genes (B and C) and downregulated genes (D and E) determined with factor loadings in PC7 were analyzed by IPA software. Dot plots show upstream regulators (B and D) or diseases and functions (C and E). (F–I) Ovaries were collected from YNG WT and *Akr1b7^−/−^* mice at 24 h after PMSG/hCG injection. (F) The immunofluorescence images show stainings with Nuclei-DAPI (blue), AKT1-TRITC (green), and pAKT1-Cy5 (red). Asterisks indicate a growing follicle belonging to the secondary or antral stage. n = 4, Scale bars, 50 µm. (G) Violin plots show the ratio of pAKT1/AKT1. n = 4, ***P < 0.001; two-tailed Student’s t test. (H) Images show DAB staining of AKT1 and pAKT1 in ovaries. Red arrow indicates a primordial follicle. n = 4, Scale bars, 20 µm. (I) Immunofluorescence images show stainings with Nuclei-DAPI (blue), KITL-TRITC (green), and AKR1B7-Cy5 (red). White arrow indicates a primordial follicle. Scale bars, 40 µm. n = 4.

### Extension of estrous cycles in *Akr1b7^−/−^* mice

Network analysis in *Akr1b7^−/−^* mice suggested that NR5A1 directly suppressed *Cyp17a1* expression, participating in progesterone metabolism in the network, including AKR1B7 (Figure 7A); LH and CG stimulation enhanced the expression of genes associated with ovulation, such as EREG and AREG (Figure 7 B). Immunohistochemical observations demonstrated that NR5A1 was present in the theca cell layer around developing follicles, and similar staining was detected in *Akr1b7^−/−^* mice (Figure 7C and 7D). The expressions of NR5A1-target genes significantly decreased in the ovaries of *Akr1b7^−/−^*mice (Figure 7E); thus, AKR1B7 might be involved in the activation of NR5A1 without affecting its expression. The staining of CYP17A1, an NR5A1-target gene, was not detected in the thecal cell layers of the ovary in *Akr1b7^−/−^* mice in contrast with WT mice (Figure 7F, 7G and Figure S5F).

**Figure 7.**
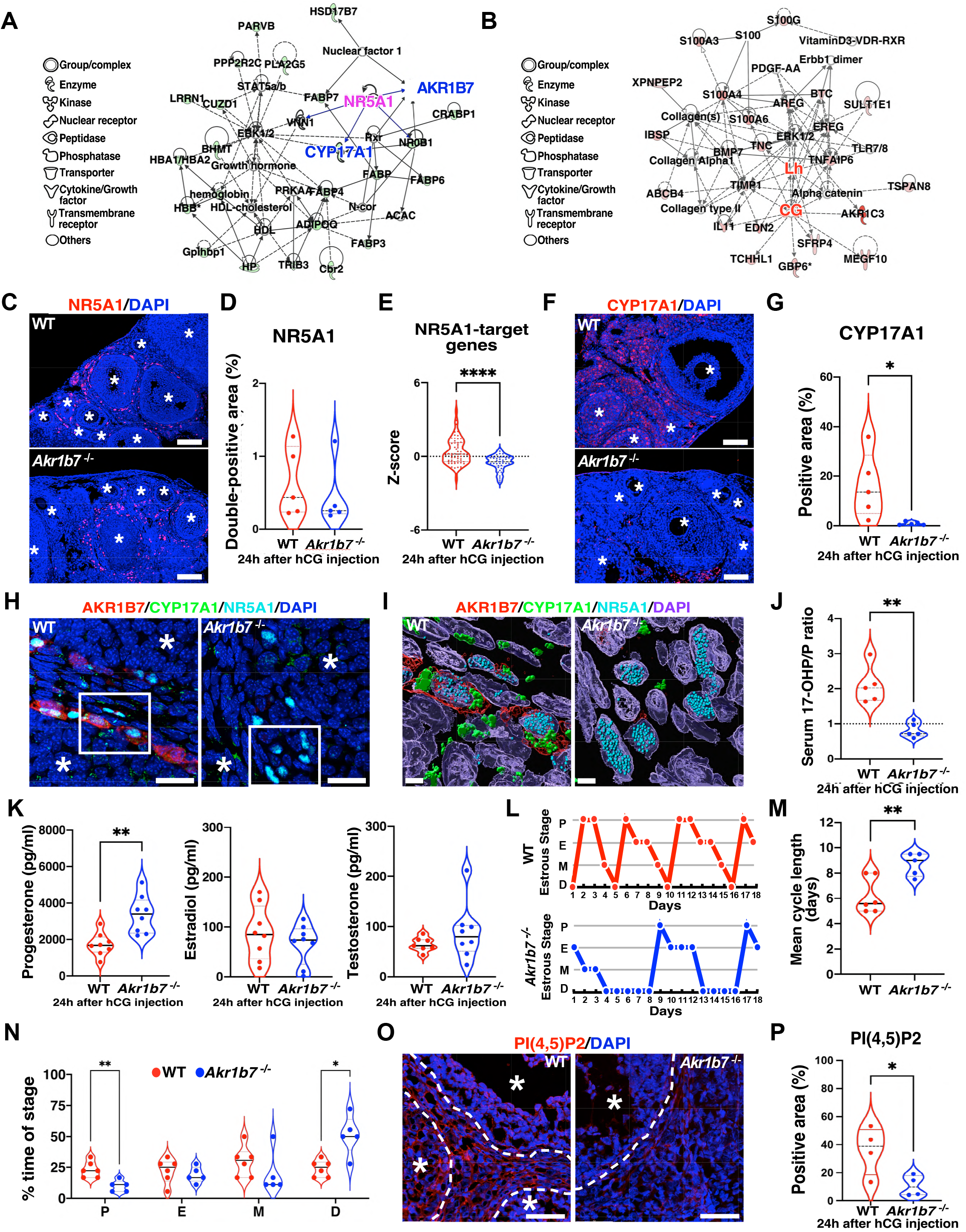
Signal pathway in thecal cell layers in *Akr1b7*-deficient mice after ovulation stimulation. (A and B) Networks show the pathways analyzed using the downregulated genes (A) and the upregulated genes (B) determined with factor loadings in PC7. (C) Immunofluorescence images show NR5A1 staining (red) in YNG WT and *Akr1b7^−/−^*ovaries at 24 h after PMSG/hCG injection. Asterisks point to a growing follicle at the secondary or antral stage. Scale bars, 100 μm. Nuclei-DAPI (blue). n = 5. (D) Violin plot shows nuclear NR5A1-positive area (%). (E) Violin plots show z-scores of RNA-seq analysis in downstream genes (*Cyp17a1, Cyp21a1*, *Cyp26b1*, *Lhcgr*, *Nr0b1*, *Vnn1*) of NR5A1. WT; n = 10, KO; n = 7. (F) Immunofluorescence images show CYP17A1 staining (red) in YNG WT and *Akr1b7^−/−^* ovaries at 24 h after PMSG/hCG injection. Asterisks point to a growing follicle at the secondary or antral stage. Scale bars, 100 μm. Nuclei-DAPI (blue). n = 5. (G) Violin plot shows CYP17A1-positive area (%). (H) Images show stainings of Nuclei-DAPI (blue), CYP17A1-FITC (green), NR5A1-TRITC (cyan), and AKR1B7-Cy5 (red) obtained using the confocal microscope. Asterisks indicate a growing follicle belonging to the secondary or antral stage. Scale bars, 20 µm. White squared area indicates theca cell layers. (I) Immunofluorescence images were processed stainings of Nuclei-DAPI (blue), CYP17A1-FITC (green), NR5A1-TRITC (cyan), and AKR1B7-Cy5 (red) to 3D images using an Imaris software. Scale bars, 4 µm. (J) Violin plots show 17-OHP/progesterone ratio in the serum of YNG WT and *Akr1b7^−/−^* mice at 24 h after PMSG/hCG injection. n = 5, **P < 0.01; two-tailed Student’s t test. (K) Violin plots show serum progesterone, estradiol, and testosterone concentrations quantified by ELISA in YNG WT and *Akr1b7^−/−^* mice at 24 h after PMSG/hCG injection. n = 8, **P < 0.01; two-tailed Student’s t test. (L) Representative patterns of estrous cycle determined by vaginal cytology are shown in WT and *Akr1b7*^−/−^ mice. Proestrus; P, estrus; E, metestrus; M and diestrus; D, n *=* 5–6. (M) Mean period length of the estrous cycle is shown using WT and *Akr1b7*^−/−^ mice. **P < 0.01; two-tailed Student’s t test. (N) Violin plots show percentages of the spending time in each stage during the estrous cycle in WT and *Akr1b7*^−/−^ mice. *P < 0.05, **P < 0.01; two-way ANOVA followed by Tukey’s multiple comparisons test. (O) Immunofluorescence images show PIP2 stainings (red) in YNG WT and *Akr1b7^−/−^* ovaries at 24 h after PMSG/hCG injection. Asterisks indicate growing follicles at the secondary or antral stage. Dotted lines represent the boundary between the theca and granulosa layer. Scale bars; 50 μm, Nuclei-DAPI (blue) (P) Violin plots show PIP2-positive area (%) in the total area of an ovarian section in Figure 7O (n *=* 4). *P < 0.05; two-tailed Student’s t test.

CYP17A1 co-localized with AKR1B and NR5A1 in the theca cell layers of developing follicles (Figure 7H and 7I). The serum levels of progesterone and its metabolites were analyzed because CYP17A1 metabolizes progesterone to 17-hydroxyprogesterone (17-OHP). Serum progesterone levels significantly increased in *Akr1b7^−/−^* mice with a reduction in the 17-OHP/progesterone ratio (Figure 7J and 7K). The estradiol and testosterone levels were not different between WT and *Akr1b7^−/−^* mice (Figure 7K). Consistent with the increase in progesterone levels, the length of the estrous cycle was significantly stretched to 9.5 d in *Akr1b7^−/−^* mice (Figure 7L and 7K); this prolongation was mainly caused by the stretching of the diestrus stage during the estrous cycle (Figure 7N). NR5A1, an upstream molecule of CYP17A1, was reported to be activated by phosphatidyl inositol 2-phosphate (PIP2),^21^ and PIP2 produced in the theca cell layers of developing follicles and stromal cells was absent in *Akr1b7^−/−^* mice (Figure 7O, 7P and Figure S5F). Thus, AKR1B7 regulates the estrous cycle through NR5A1/CYP17A1/ progesterone, which might be activated by PIP2 production.

During ovarian aging, the transient expression of *Akr1b7* observed in the theca cell layers of young mice was attenuated independent of ovulation stimulation. Consistent with AKR1B7 expression, KITL staining was weakly detected only in stromal cells of aged ovaries, distinct from the apparent staining of young ovaries (Figure S6A and S6B). Akt activation during ovarian aging was almost undetectable in the whole ovary, including in the oocytes of primordial follicles, despite Akt expression (Figure S6C and S6D). CYP17A1 detected in young ovaries administrated with CG almost disappeared around the theca cell layers of aged ovaries (Figure S6E and S6F). The ovarian progesterone level was also significantly increased according to the reduction of 17-OHP, a metabolite of progesterone, by CYP17A1 according to ovarian aging, reflected in serum progesterone and 17OHP levels (Figure S6G, S6H, and Figure S7). Consistent with sustained progesterone levels, the estrous cycle was extended according to the prolonged diestrus stage (Figure S6I–K). The signaling pathways attenuated in the aged mice were consistent with those in the *Akr1b7^−/−^* mice.

## Discussion

This study comprehensively analyzed mRNA expression during follicular development and ovarian aging. In response to ovulation stimulation, the aldo-keto reductase AKR1B7 pathway activated in young mice markedly diminished in the old mice. *Akr1b7^−/−^* mice without the start codon enhanced ovulation of immature oocytes with a decline in primordial follicles. Oocyte Akt activation in primordial follicles, essential for follicular development, was suppressed in *Akr1b7^−/−^* mice, consistent with decreased KITL expression in *Akr1b7*-expressed stromal cells. *Akr1b7^−/−^* mice simultaneously exhibited a prolonged estrous cycle via the sustained progesterone level due to the reduced CYP17A1 in the thecal cell layers expressing *Akr1b7*.

A decline in the quantity and quality of oocytes is a typical characteristic of ovarian aging. They are especially associated with the developmental failure of follicles in the early phases, such as primordial follicles. Ovarian aging manifests as reproductive dysfunction, such as an extended menstrual cycle and hormonal abnormalities.^2^ In the ovarian aging mouse model used in this study, the number of primordial to secondary follicles significantly decreased during the estrous cycle. The marker genes in each follicle based on scRNA-seq showed reduced primordial and primary follicles in aging ovaries, in contrast with the upregulation of marker genes in secondary follicles. Pathway analysis demonstrated that the LH/CG-regulated AKR1B7 pathway was reduced, and the expression of genes related to glucose and lipid metabolism was downregulated in the ovarian aging mouse model. Ge et al. reported that AKR1B7 regulated lipid and glucose homeostasis in the liver,^22^ indicating that decreased *Akr1b7* expression may affect downstream pathways such as glucose and lipid metabolisms. *Akr1b7* is predominantly expressed in the stromal and theca cell layers of young mice, and its expression continues at low levels during the estrous cycle in aging ovaries. However, the ICA and 4-HNE activities of AKR1B7 remained slightly reduced during ovarian aging, in contrast with their expression. We speculated that AKR1B7 might be implicated in the function(s) except for detoxification in the ovary. *Akr1b7^−/−^* mice produced using i-GONAD showed accelerated ovulation of immature oocytes with relatively small diameters, leading to decreased primordial follicles and increased CLs according to the age of the mice. RNA-seq analysis using the whole ovary of the *Akr1b7^−/−^*mice indicated that disorders of glucose metabolism and diabetes were found in “Diseases and Functions.”

The ovulation of immature oocytes observed in *Akr1b7^−/−^*ovaries may be mediated by Akt activation for the following reasons. First, Akt activation, essential for follicular development in primordial follicles in response to FSH,^23^ was absent in the oocytes of primordial follicles in *Akr1b7^−/−^* mice. Second, *Akt^−/−^* ovaries treated with exogenous gonadotropins induced reduced primordial follicles and increased secondary follicles^19^ and caused a delay in the onset of the estrous cycle,^24^ which were almost consistent with those of *Akr1b7^−/−^* mice. Finally, Akt inactivation is associated with glucose metabolism disorders related to diabetes, as detected in *Akr1b7^−/−^* mice.^25^

How does AKR1B7 present in stromal cells activate oocyte Akt in primordial follicles? Human AKR1B10, an ortholog of mouse AKR1B7 (Alliance of Genome Resources, Apr 2022), is bound to acetyl-CoA carboxylase (ACC), a rate-limiting enzyme in fatty acid synthesis that prevents the degradation of ACC.^26^ AKR1B7 deletion may cause the degradation of ACC due to the dissociation, as well as AKR1B10. As shown in Figure 6E, the pathway analysis of *Akr1b7^−/−^*mouse ovaries revealed the decreased concentrations of lipids and triacylglycerol as a downregulated pathway, indicating reduced lipid synthesis. The reduced ACC results in a metabolic switch to a glycolysis system due to the inhibition of the conversion of acetyl-CoA to malonyl-CoA.^27^ The prevention of mitochondrial respiration induces an increase in cellular AMP/ADP levels, which causes AMPK activation.^28^ AMPK activation may cause Akt inactivation through decreased KITL because AMPK inhibition triggers KITL release, causing oocyte Akt activation in primordial follicles.^29^ In the ovary, KITL staining was detected in the cytoplasm of *Akr1b7*-expressed stromal cells, in addition to the membrane staining of granulosa cells, indicating that *Akr1b7* may regulate cytoplasmic KITL2 expression, distinct from the membrane KITL1 of granulosa cells. Consistent with our data, human KITL expression was observed in granulosa and stromal cells.^30^ The introduced KITL restored fertility in fertile mice.^31^ Reduced KITL expression in the stromal cells of *Akr1b7^−/−^* mice might be associated with primordial follicle arrest.

The network pathways activated according to the time course after ovulation stimulation showed differences between young and old mice. The ovaries of young mice showed that estrogen and its receptors activated the gonadotropin-releasing hormones *Gnrh2* and *Pgh2*, associated with hCG activation 48 h after ovulation stimulation.^32,33^ At 72 h, CG activation might have been involved in the induction of growth hormones, such as *Vegf* and *Igfbp4I*, in the early phase. After activation, the CG, FSH, and LH pathways showed the induction of the expression of metabolic enzymes for sex hormones such as *Cyp17a1* and *Hsd17b3*.^34,35^ In the aged ovary, the major pathways activated by sex hormones, except for exogenous LH and FSH, were not detected until 96 h after stimulation. Scaffold proteins, such as Shank1, Hsp70, Hsp90, proteinase, caspase, and 26S proteasome, were detected as the activated pathways, indicating that protein denaturation and degradation occurred during the cycle. Protein denaturation may activate immune cells because gene markers related to macrophages and T cells, such as NADPH oxidase, IL4, TCR, and CD3, are present in these pathways. Reports have indicated that inflammation and oxidative stress are closely associated with ovarian aging.^13,14,36^ During the formation of the secondary follicles, theca cells are recruited by the differentiation from the progenitor cells in stromal cells.^37^ The marker genes of theca cells, *Cyp17a1, Col1a2*, and *InsI3* mRNA, were significantly decreased in old mice,^38^ indicating that the differentiation of stromal cells into theca cells may be defective in aged ovaries. In a previous report, theca cells participate in steroidogenesis, which is necessary for folliculogenesis and the estrous cycle.^39^ As shown in this study, *Akr1b7* was localized in theca cells and stromal cells. According to the literature, the major activities of AKR1B7 are the detoxification of steroid metabolites and the formation of PGF_2α_.^18,40^ The activities of toxic metabolites were observed in *Akr1b7^−/−^* ovaries, similar to those in WT mice, although the activities were reduced in old mice with decreased expression of *Akr1b7*. Part of the AKR1B7 activities of *Akr1b7^−/−^*mice may be compensated by AKR1B8 with high homology to *Akr1b7* because *Akr1b8* mRNA was maintained in *Akr1b7^−/−^*mice, in contrast with the reduction of *Akr1b8* in old mice. AKR1B8 might have difficulty compensating for all AKR1B7 activities in the theca cells because AKR1B8 has distinct from the substrate specificity from AKR1B7 expression, except for ICA and 4-HNE.^41^ The metabolic pathways implicated in sex hormones are activated in the theca cells, and sex hormones strictly regulate the estrous cycle.^2^ Compared to WT mice, the estrous cycle was prolonged in the Akr1b7^−/−^ mice. The progesterone level lasted at a high level, coincident with a previous report using mice deficient in exons 2-4 of Akr1b7.^42^ The increase in the progesterone level was accompanied by a reduced 17-OH-metabolite, which was consistent with the decrease in the expression of *Cyp17a1* localized in theca cells. *Cyp17a1* expression is regulated by NR5A1/SF-1,^43^ the target genes of which showed decreased expression in *Akr1b7^−/−^*mice, despite expression and localization of NR5A1/SF-1 remaining unchanged, suggesting that the ligand is suppressed in KO mice. NR5A1/SF-1 directly associates with PIP2, which is converted to PIP3, and functions as a ligand for NR5A1/SF-1.^44,45^ Exposure to gonadotropins led to PIP2 formation in the theca cell layers and the boundary regions of granulosa cell layers; the PIP2 product was eliminated in KO mice as shown in Figure 7O. In the breast cancer cell line-introduced siRNA of *Akr1b10*, a human ortholog of *Akr1b7*, the PIP2 product is significantly decreased, reducing cell growth. Several studies have implicated phospholipid metabolism in the AKR superfamily. AKR1B10 regulates the stability of acetyl-Co carboxylase, which may indirectly contribute to the synthesis of membrane lipids, such as PIP2.^46^ Spite et al. reported that the AKR1B superfamily has an enzymatic activity against phospholipid aldehydes present in the PIP2 structure,^47^ which may be directly metabolized by AKR1B7 as a substrate. The reduction in phospholipids by AKRs could regulate the signaling mechanisms triggered by biological phospholipids, including the aldehyde structure.^48^ PIP2 generated by AKR1B7 controls the estrous cycle through progesterone/CYP17A1/NR5A1 and may also affect the normal development of primordial follicles through Akt activation.

In conclusion, *Akr1b7*, present in the theca and stromal cells of the ovary, induces KITL expression in response to ovulation stimulation, leading to the activation of follicular development through Akt activation. Furthermore, *Akr1b7* controls the estrous cycle through progesterone metabolism via *Cyp17a1* expression. During aging, the decreased *Akr1b7* accelerates the ovulation of immature oocytes and prolongs the estrous cycle, as is frequently observed in ovarian aging. Regulation of *Akr1b7* expression would be an essential role in increasing the fertility rate during ovarian aging.

## Acknowledgments

This work was the result of using research equipment shared in MEXT Project for promoting public utilization of advanced research infrastructure (Program for supporting construction of core facilities) Grant Number JPMXS0440400021-23 and grants-in-aid for scientific research (Grant Nos. 20K17491, 21K08644, 21H03358, and 22K19711) from of the Ministry of Education, Culture, Sports, Science and Technology of Japan.

We thank Ms. Kaori Kaminoyama (Center for Plant Sciences, Kyoto Sangyo University) for the next-generation sequencing (NGS) analysis using NextSeq500. We also thank Ms. Makiko Nakagawa, Ms. Takako Moriyama, and Ms. Yuko Nakatani (Yamaguchi University) for their technical support in NGS analysis and the technical support at the Yamaguchi University Science Research Center.

## Author Contribution

Conceptualization, K.I. and Y.M.; Methodology, K.I., K.W., M.O., S.K., T.H., S.S., and Y.M.; Investigation, K.I. and K.W.; Writing, K.I. and Y.M.; Funding Acquisition, K.I., K.W., and Y.M.; Supervision, T.M., M.A., H.T., N.S., and Y.M.

---

**Figure S1 Correlation of mRNA expression level between whole transcriptome analysis (WTA) and qPCR**

(A) mRNA expression in WTA with superSAGE (counts per million) and qPCR (relative value using *Gapdh* as a control) were plotted to examine the correlation. Ratios (48 h/24 h, 72 h/24 h, and 96 h/24 h) of mRNA levels in the indicated genes were log_2_ transformed and plotted. n = 3, Correlations were valued with the Pearson correlation coefficient (r). (B) Violin plots showed that time course mRNA expressions the 4 genes measured by WTA with superSAGE (red) and qPCR (blue) in YNG and OLD mice. The relative expression levels at each time point were normalized to the level at 24 h.

**Figure S2 mRNA expression levels of aldo-keto reductase superfamily in ovarian aging**

mRNA expression levels of the aldo-keto reductase superfamily in YNG and OLD ovaries were analyzed using the superSAGE method at the indicated times after PMSG/hCG injection. Expression levels are displayed as CPM. n = 3. Two-way ANOVA followed by Tukey’s multiple comparison test; *P < 0.05, ***P<0.001.

**Figure S3 IPA network analysis during estrous cycle after ovulation stimulation in ovarian aging**

Volcano plots indicate upregulated (red) or downregulated genes (green) with a 1.5-fold change having P value < 0.05 at the indicated times in YNG (A–C) and OLD (D–F). The regulated genes at each time point were analyzed using IPA network analysis, and each first network is shown.

**Figure S4 Gene expressions of off-target sequences in genome-editing mouse of *Akr1b7***

(A) Off-target sequences were analyzed with software, CRISPR-Cas9 gRNA checker (Integrated DNA Technologies), and the top 10 off-target regions using the guide RNA targeting of *Akr1b7* were shown. A lower score indicates a higher risk of genome editing. (B) Mapping of RNA-seq reads is shown on the whole exon (upper) and exon-1(lower) in *Akr1b3*, *Akr1b8*, and *Akr1b10*. Colored reads indicate single reads mapped in the forward direction (green), in the reverse direction (red), and paired reads (blue). Yellow reads indicate nonspecific mapping that has identical sequences to other genes.

**Figure S5 Confirmation of factors regulated by AKR1B7 using *Akr1b7^−/−^* mice ovary**

(A) Sections were prepared from WT and *Akr1b7^−/−^* YNG mice ovaries at 24 h after 5 IU PMSG/hCG injection and were stained with the indicated antibodies and DAPI. Circles with dotted lines show the oocyte of primordial follicle. Max intensity projections of z-stack images in 4 μm section were obtained with a 100x objective lens and 3.0x digital zoom using a confocal microscope. Nuclei-DAPI and pAKT1-Cy5 were observed by excitation at 405 and 638 nm, respectively. Scale bars, 20 μm. (B) Violin plots show pAKT1 intensities in the oocyte of the primordial follicle. n = 17–18, ****P < 0.0001; two-tailed Student’s t test. (C) Ovary extracts collected from WT and *Akr1b7^−/−^* YNG mice ovaries at 24 h after 5 IU PMSG/hCG injection were subjected to immunoblotting using anti-AKT1 antibody and anti-pAKT1 antibody. n = 4–5. (D) Violin plots show the ratio of band intensities of pAKT1 and AKT1 in Figure S5C. *P < 0.05; two-tailed Student’s t test. (E) Sections were prepared from 16-week-old WT and *Akr1b7^−/−^* mice ovaries at 24 h after 5 IU PMSG/hCG injection and were stained with the indicated antibodies and DAPI. Arrows show the primordial follicle. Scale bars, 20 μm. n = 4. (F) Sections were prepared from ovaries of WT and *Akr1b7^−/−^* YNG mice at 24 h after 5 IU PMSG/hCG injection and were stained with anti-AKR1B, anti-NR5A1, or anti-PI(4,5)P2 antibodies and DAB reagent. Sections using anti-CYP17A1 antibodies were stained purple. n = 4–5.

**Figure S6 Expressions of factors regulated by AKR1B7 in ovarian aging**

(A) Sections were prepared from ovaries of YNG and OLD mice at 24 h after 5 IU PMSG/hCG injection and were stained with anti-AKR1B7 (Cy5, red) and anti-KITL (TRITC, green) antibodies and DAPI (blue). Arrows point to a primordial follicle. Scale bars, 20 μm. n = 5. (B) Violin plots show the positive areas (%) of AKR1B or KITL on tiling images of the whole ovary. n = 5, *P < 0.05; two-tailed Student’s t test. (C) Sections were prepared as described in Figure S6A and stained with anti-pAKT (Cy5, red) and anti-AKT antibodies (TRITC, green) and DAPI (blue). Circles with dotted lines represent the oocyte of primordial follicle. Scale bars, 20 μm. (D) Violin plots show the ratio of fluorescent intensities of pAKT1 and AKT in the region as shown in Figure S7C. n = 7–11, *P < 0.05; two-tailed Student’s t test. (E) Sections were prepared from ovaries of YNG and OLD mice at 96 h after 5 IU PMSG/hCG injection and stained with anti-CYP17A1 antibody (Cy5, red) and DAB reagents. Scale bars, 50 µm. (F) Violin plots show the positive areas (%) in Figure S6E. n = 3–4, **P < 0.01; two-tailed Student’s t test. (G and H) Progesterone and 17α-OHP levels and the ratio of 17α-OHP to progesterone were quantified using the reversed-phase HPLC method. The violin plot shows progesterone and the metabolite levels in the ovary (G) or the serum (H) obtained from YNG and OLD female mice at 96 h after 5 IU PMSG/hCG injection. n = 3–4, *P < 0.05, **P < 0.01, ***P < 0.001; two-tailed Student’s t test. (I) Representative pattern of the estrous cycle determined by vaginal cytology is shown in YNG and OLD mice. Proestrus; P, estrus; E, metestrus; M and diestrus; D, n *=* 5. (J) Mean period length of the estrous cycle determined from Figure S6I is shown in YNG and OLD mice. *P < 0.05; two-tailed Student’s t test. (K) Violin plots show percentages of the spending time in each stage during the estrous cycle determined from Figure S6I in YNG and OLD mice. *P < 0.05, **P < 0.01; two-way ANOVA followed by Tukey’s multiple comparisons test.

**Figure S7 mRNA expressions in genes associated with progesterone metabolism during estrous cycle after ovulation stimulation**

(A) Progesterone metabolism. A steroidogenic acute regulatory protein (StAR) regulates cholesterol transfer within the mitochondria. In the mitochondria, the enzyme P450 side-change cleavage (CYP11A1, also known as P450scc) converts cholesterol into pregnenolone, which is further transformed into progesterone in the endoplasmic reticulum by 3*β*-hydroxysteroid dehydrogenase (HSD3B1). Progesterone is metabolized by P450c17 (CYP17A1), 20α-hydroxysteroid dehydrogenase (AKR1C18, also known as 20α-HSD), 3-oxo-5α-steroid 4-dehydrogenase (SRD5A1, 2, and 3), and/or 3α-hydroxysteroid dehydrogenase (AKR1C14, also known as 3α-HSD). (B) mRNA expression levels of *Star, Cyp11a1, Hsd3b1, Cyp17a1, Akr1c18, Akr1c14, Srd5a1, Srd5a2,* and *Srd5a3* in YNG and OLD ovaries at the indicated times after 5 IU PMSG/hCG injection. n = 3. Two-way ANOVA followed by Tukey’s multiple comparison test; *P < 0.05.

## STAR★Methods

### Resource availability

#### Lead contact

Further information and requests for resources and reagents should be directed to and fulfilled by the lead contact, Yoichi Mizukami.

### Materials availability

Generated mouse line in this paper will be made available on request.

#### Experimental models and subject details

##### Animals and the collection of ovaries

Female C57BL/6N mice at the age of 6–10 weeks (“YNG”) and 48–56 weeks (“OLD”) were purchased from Japan SLC Inc (Shizuoka, Japan). To synchronize the estrous cycle, we performed a superovulation procedure by administering pregnant mare serum gonadotropin (PMSG, 5 IU; ASKA Pharmaceutical, Tokyo, Japan) and human chorionic gonadotropin (hCG, 5 IU; ASKA Pharmaceutical) at 48 h intervals to mice. Mice were euthanized 24, 48, 72, and 96 h after hCG administration. Ovaries were collected at indicated time points. One side of the ovary was quickly frozen in liquid nitrogen and stored in RNAlater (Thermo Fisher Scientific, Waltham, MA) at −80°C for the expression analyses.

The other side was fixed with 4% paraformaldehyde and embedded in paraffin for histological analysis. ICR mice (Japan SLC Inc.) were used to generate gene-edited mice and maintained in an ICR genetic background. The primers and TaqMan probes used for genotyping are listed in Table S1. Females were group-housed with up to 5 mice per cage under a stable temperature in a 12 h light and dark cycle with water and food *ad libitum*. All experiments were performed in accordance with the recommendations of the Guide for Animal Experiments at the Yamaguchi University School of Medicine. The Committee on the Ethics of Animal Experiments of Yamaguchi University School of Medicine reviewed and approved all procedures.

##### Histochemical analysis

Pretreatment of the tissues was performed as previously described.^1^ The fixed ovaries with 4% paraformaldehyde were embedded in paraffin block and serially sectioned at 8 μm using a microtome. The sections were stained with hematoxylin and eosin (H&E). Classification and counting of ovarian follicles at different stages were performed as previously described.^2^

Follicles were classified as follows: the follicles with oocyte surrounded by a single layer of squamous granulosa cells were characterized as primordial follicles, the follicles with oocyte surrounded by a single layer of cuboidal granulosa cells were characterized as primary follicles, the follicles with oocyte surrounded by multiple layers of cuboidal granulosa cells with or without antral space development were characterized as secondary follicles, and the follicles with oocyte situated on cumulus oophorus with multiple layers of granulosa cells and a large confluent antral space were characterized as antral follicles.

The total number of primordial and primary follicles was counted in every fifth section of the ovary. Because the minimum diameters for primordial and primary follicles were 7–25 μm ^3^, smaller than 40 μm, corresponding to an 8 μm x fifth section interval, raw counts were multiplied by 5 to obtain the estimated number. The total number of secondary and antral follicles was counted using the raw number for each fifth section. Repetitive counting was avoided by counting only follicles containing an oocyte with a visible nucleolus. The number of CLs, including ovulated luteinizing follicles, was counted using the raw number to avoid repetition among serial sections.

##### Laser capture microdissection

Ovaries were embedded in paraffin block, and 12 μm sections were mounted on membrane glass slides (Thermo Fisher Scientific) coated with 0.1% poly-L-lysine solution in H_2_O (Sigma-Aldrich, St. Louis, MO). Slides were deparaffinized with xylene, stained with H&E, and loaded onto the stage of a laser microdissection apparatus (LMD6500; Leica microsystems, Wetzlar, Germany). The granulosa or theca cell layers were captured using an infrared capture laser under a microscope, and approximately 50 layers per sample were collected. The RNeasy formalin-fixed paraffin-embedded kit (Qiagen, Hilden, Germany) was used to extract total RNA.

##### Total RNA isolation and quantitative polymerase chain reaction analysis (qPCR)

qPCR was performed as previously described.^4,5^ In brief, the RNeasy Mini Kit was used to isolate total RNAs from each ovary according to the manufacturer’s protocol. After the reverse transcription procedure, messenger RNA (mRNA) expression values were measured using the QuantiTect SYBER Green polymerase chain reaction (PCR) kit (Qiagen) with a Rotor-Gene 6000 (Qiagen) or CFX384 (Bio-Rad, Hercules, CA). Primers used for qPCR are listed in Table S1. The relative quantity of each gene was normalized to that of *Gapdh* or *Actb* gene expression.

##### Whole transcriptome analysis (WTA) by serial analysis of gene expression (SAGE) method

Total RNA was extracted from the ovaries of mice by using the RNeasy Mini Kit (Qiagen). The SOLiD™ SAGE™ Kit barcoding adapter module (Thermo Fisher Scientific) was used to construct libraries of 27-bp tags adjacent to the 3′ends of mRNAs according to the manufacturer’s protocol. Each library was amplified by PCR with a different barcoding primer using the SOLiD RNA Barcoding Kit and purified using the Pure-Link PCR Micro Kit (Thermo Fisher Scientific). The cDNA concentration of each library was measured by qPCR using a Rotor-Gene 6000 (Qiagen) and adjusted to 500 pM for optimal input into the emulsion PCR. Emulsion PCR for the enrichment of libraries was performed according to the protocol of SOLiD™ EZ Bead System (Thermo Fisher Scientific) as described previously.^6^ SOLiD5500 (Thermo Fisher Scientific) was used to sequence libraries of 27-bp tags. CLC genomics workbench (Qiagen) was used to trim the sequencing reads and align them to a reference mouse genome (GRCm38), as previously described.^1^ Reads mapped to exon regions of each gene were counted. mRNA abundance was expressed as tag counts per million (CPM). Differentially expressed genes were defined as those with an absolute value of fold change >1.5 and normal P < 0.05 between a pair of samples. A volcano plot plotting the value of log_2_ (fold change) against -log_10_ (P value) was created using R version 3.3.0. Gene expression differences with the largest variance were visualized with a heatmap using JMP pro14 (SAS Institute Japan, Tokyo, Japan). To validate the intragroup homogeneity among age and time points, principal component analysis (PCA) was performed using JMP pro14. Network speculation and functional annotation of differentially expressed genes were performed using ingenuity pathway analysis (IPA) (Qiagen).

##### WTA by RNA-seq analysis

RNA-seq analysis was performed as previously described.^7-11^ The RNeasy Mini Kit (Qiagen) was used to extract total RNA from the ovaries of mice, and mRNA was purified using Oligo dT beads (NEBNext Poly (A) mRNA magnet Isolation Module, New England Biolabs, NEB). The NEBNext Ultra II RNA Library Prep Kit (NEB) and NEBNext Multiplex Oligos for Illumina (NEB) were used to generate cDNA libraries for Illumina sequencing. An Agilent 2200 TapeStation (D1000, Agilent, Santa Clara, CA) was used to evaluate the quality and concentration of the libraries. The confirmed libraries were mixed in equal molecular amounts for clustering and sequencing on an Illumina Next-seq500 DNA sequencer with a 75 bp paired-end cycle sequencing kit (Illumina, San Diego, CA). Trimmed reads were mapped to the mouse reference genome GRCm38 release-92 using the default settings. For pathway analysis, factor loadings were calculated with the PCA of the JMP Pro version 14.0 software, the top 200 genes were selected, and the pathway analysis for the detected genes was examined using IPA (Qiagen).

##### Measurement of enzymatic activity

The cytosol extracts of ovaries in 0.1 M sodium phosphate buffer (pH 6.6) containing 1 mM dithiothreitol and protease inhibitor cocktail (Sigma-Aldrich) were collected by centrifugation at 105,000 x g for 60 min at 4°C. The standard reaction mixture consisted of 0.1 M sodium phosphate buffer (pH 6.6), 1 mM isocaproaldehyde (Santa Cruz Biotechnology, Dallas, TX), or 640 μM 4-hydoroxynonenal (Santa Cruz Biotechnology) as a substrate, and 1 mM NADH (Sigma-Aldrich). The reaction mixture was prewarmed at 25°C for 10 min in a 96-well plate (#Clear black plate; Greiner Bio-One, Kremsmünster, Austria), and the reaction was initiated by adding cytosolic extract. The reductase activity for each substrate was determined by the consumption rate of NADH at 340 nm for 20 min, with 405 nm as a reference, using a FlexStation 3 microplate reader (Molecular Devices, San Jose, CA). Enzymatic characterization was performed using Lineweaver-Burk plots as described previously.^12,13^ Values of the reductase activity were normalized by the total protein concentration of cytosol extracts determined using the method of Bradford (Bio-Rad).

##### Measurements of PGF_2α_

Whole ovaries were homogenized with a chilled bead mill at 3,200 rpm for 30 s in 0.1 M phosphate buffer (pH 7.4) containing 1 mM EDTA and 10 μM indomethacin. A PGF_2α_ enzyme immunoassay kit (Cayman Chemical, Ann Arbor, MI) was used to measure the concentrations of PGF_2α_ in the supernatants according to the manufacturer’s instructions.^14^ A FlexStation 3 Micro plate reader (Molecular devices) was used to measure the absorbance at a wavelength of 405 nm. The total protein content of the supernatant was determined using the method of Bradford (Bio-Rad).

##### Immunohistochemical staining

Ovaries were fixed in 4% paraformaldehyde at 4°C overnight. The tissues were embedded in paraffin blocks and cut into 4 μm thick tissue sections. Sections were immunostained, reacted with 3,3′-diaminobenzidine (DAB) chromogen, and counterstained with hematoxylin as previously described.^1,15^ Primary antibodies used for immunohistochemistry were anti-human AKR1B10 rabbit polyclonal antibody with an identical epitope sequence against mouse AKR1B7 (Thermo Fisher Scientific, #PA5-22036, 1:1500) and anti-DNA/RNA damage mouse monoclonal antibody (Abcam, Cambridge, UK, ab62623, 1:2000). Quantitative analysis of AKR1B7 expression in sections was performed using digital images. The resulting images were divided into individual DAB and hematoxylin images by using a color deconvolution plugin in ImageJ (National Institutes of Health). The threshold parameters were set at 130 Gy for DAB staining and 200 Gy for hematoxylin staining. The trimmed DAB and hematoxylin images were transformed to pseudocolored red and blue, respectively, and overlaid using MetaMorph imaging software (Molecular Devices). The intensity of DAB staining (red) was measured in the theca layer of each follicle, and the DAB intensity per area was calculated. The theca layer was defined as a layer of elongated cells approximately 3–5 cell layers thick, immediately adjacent to the basal lamina of an ovarian follicle.^16^

##### Immunofluorescence staining and 3D imaging

From 4 to 5 individual ovaries for each genotype were embedded in one paraffin block and cut into 4 μm thick tissue sections. For comparative analyses, tissue sections obtained from WT and *Akr1b7*^−/−^ ovaries were mounted on the same slide. An automatic procedure was performed using Ventana Benchmark Ultra (Roche Diagnostics, Basel, Switzerland). In brief, the sections were deparaffinized and heated at 95°C for 64 min with antigen retrieval buffer, Tris-EDTA buffer (CC1; Roche Diagnostics), or citrate buffer (CC2; Roche Diagnostics). The 1st antibody buffer containing 5% serum and 0.1% Triton X was hand-applied to the drops on the slides and incubated at 37°C for 32 min. The dilution factor in the 1st antibody buffer and retrieval buffer in the antibodies were as follows: Anti-AKR1B10 rabbit polyclonal antibody (Thermo Fisher Scientific, PA5-22036, 1:2000, CC1), anti-CYP17A1 rabbit monoclonal antibody (Abcam, ab125022, 1:100, CC2), anti-NR5A1 rabbit monoclonal antibody (Cell signaling technology, 12800S, 1:50, CC2), anti-phosphorylated (Ser473) AKT1 rabbit monoclonal antibody (Abcam, ab81283, 1:50, CC2), anti-AKT1 rabbit monoclonal antibody (Cell signaling technology, #75692, 1:20, CC2), anti-AKT (pan) rabbit monoclonal antibody (Cell signaling technology, #4691, 1:600, CC2) and anti-KITL/SCF rabbit polyclonal antibody (Abcam, ab64677, 1:400, CC1). Omni-Map anti-rabbit horseradish peroxidase (HRP; Roche Diagnostics) was incubated at 37°C for 16 min and then reacted with FITC, TRITC, or Cy5 fluorophores. Sections were coverslipped using VECTASHIELD with DAPI or a Vector TrueVIEW Autofluorescence Quenching Kit with DAPI (Vector Laboratories, Burlingame, CA). Fluorescent images were captured with a z-stack of a 4 μm thick section by using a lightning mode of STELLARIS STED confocal microscope (Leica microsystems), and 3D images were generated using Imaris software (Oxford instruments, Oxford, UK). In 3D rendering using Imaris, Nuclei-DAPI, AKR1B-Cy5, and CYP17A1-FITC signals were processed in the 3D surface mode, and NR5A1-TRITC was processed with a 3D spot object. Images were corrected at both ends of the luminance histogram. For comparative analysis, the acquisition, correction, and 3D creation settings were identical for the WT and *Akr1b7*^−/−^ images.

##### Immunofluorescence staining of PI(4,5)P2

The ovaries were frozen in Tissue-Tek O.C.T. compound (Sakura Finetek, Tokyo, Japan). Frozen sections were cut 8 μm thick and mounted on slides as previously described.^14^ The sections were fixed with acetone at -30°C for 10 min and air-dried at RT for 5 min. The slides set using Ventana Benchmark Ultra. Goat F(ab) anti-mouse IgG H&L (Abcam) was incubated at 37°C for 16 min to block endogenous mouse IgG staining. The 1st antibody buffer containing anti-PI(4,5)P2 mouse monoclonal antibody (1:500, Echelon Biosciences, Salt Lake City, UT), 5% BSA, and 0.5% Triton X was hand-applied to the drop on the slides and reacted at 37°C for 32 min. Omni-Map anti-mouse HRP (Roche Diagnostics) was incubated at 37°C for 8 min and then reacted with Cy5 fluorophores. The sections were coverslipped using the Vector TrueVIEW Autofluorescence Quenching Kit with DAPI (Vector Laboratories, Burlingame, CA).

##### Quantitative analysis on immunofluorescence staining

Digital images were acquired using an all-in-one fluorescence microscope (BZ-X800, Keyence, Osaka, Japan) and analyzed using a hybrid cell counting tool on a BZ-X800 analyzer. For comparative analyses, the acquisition settings were identical for each protein dataset. The areas of the DAPI-, TRITC-, and Cy5-positive regions were measured using fluorescence signals above the threshold value on tiling images of the whole ovary. The total area of the ovarian sections was measured using the DAPI-positive area. The oviduct and fat pad surrounding the ovary were manually excluded using the trimming tool in the software. The positive area for the targeted protein in each section was normalized to the total area of the ovarian section.

##### Western blotting

Whole ovaries were homogenized in RIPA lysis buffer. The supernatants were collected after centrifugation. Electrophoresis and western blotting were performed after the addition of SDS sample buffer as previously described.^17-19^ The Extracts were electrophoresed in 10% (w/v) polyacrylamide gels in the presence of SDS. The gels were transferred to PVDF membranes. The membrane was blocked and incubated with anti-AKT1 rabbit monoclonal antibody (Cell Signaling Technology, Danvers, MA, cat#75692, 1:1000) or anti-phosphorylated (Ser473) AKT1 rabbit monoclonal antibody (Abcam, ab81283, 1:1000) at 4°C overnight. Anti-rabbit IgG-conjugated HRP (1:10000) was incubated for 30 min at RT. Antigens were visualized using enhanced chemiluminescence detection reagents and observed using an Amarsham Imager (Cytiva, Marlborough, MA). The densities of the detected bands were measured using ImageQuant TL software (Cytiva).

##### Generation of *Akr1b7*-deficient mice using i-GONAD-mediated genome editing

i-GONAD was performed as previously described.^20,21^ In brief, surgical procedures were performed on anesthetized ICR females on pregnant day 0.7 under observation by using a dissecting microscope (SZ60; Olympus, Tokyo, Japan). The reproductive tract (ovary-oviduct-uterus) was gently pulled out of the abdominal cavity, and the position was fixed by holding the fat pad with an artery clip. Moreover, 1.5 μL per an oviduct of Opti-MEM (Thermo Fisher Scientific) containing 30 μM annealed *Akr1b7*-crRNA/tracrRNA (Integrated DNA Technologies, Coralville, IA) and 1 mg/μL CAS9 protein (Integrated DNA Technologies) was injected into the oviductal lumen from upstream of the ampulla by using a micropipette connected to a mouthpiece (KITAZATO corp, Shizuoka, Japan). The oviductal ampulla filled with CRISPR/CAS9 solution was covered with a piece of wet paper (Kimewipe, Japan) soaked in PBS and grabbed with a tweezer-type electrode. Electroporation was performed using CUY21EDIT-II (BEX, Tokyo, Japan) under the following parameters: square (mA), Pd V:80 V, Pd A:150 mA, Pd on:5.00 ms, Pd off:50 ms, Pd N:3, Decay:10%, and Decay-Type: Log (+/−). After the surgical procedure, the mice were incubated on a heating pad at 37°C until they awoke from the anesthesia.

##### Monitoring of estrous cycle

Vaginal smears on a microscope slide were stained with Giemsa’s stain solution (Nacalai Tesque, Kyoto, Japan), as previously described,^14^ and classified into 1 of 4 estrous stages, as previously described: proestrus, estrus, metestrus, and diestrus.^22^ The mean cycle length represents the mean of the days between the estrus and the next estrous cycle.

##### Measurements of steroid hormones in serum using ELISA

Serum was separated from whole blood by centrifugation and mixed with 4x the sample volume of diethyl ether. The extracts into organic solvent were dried at 40°C under the nitrogen stream and dissolved with enzyme-linked immunosorbent assay (ELISA) buffer (Cayman Chemical). ELISA was performed for estradiol (E2), testosterone, and progesterone (Cayman Chemical), according to the manufacturer’s protocol. A FlexStation 3 microplate reader (Molecular Devices) was used to measure absorbance at a wavelength of 405 nm.

##### Measurements of progesterone metabolites in ovary and serum

The homogenate of the ovary or the serum obtained from whole blood was dissolved by adding 5x the sample volume of ethyl acetate that contained one ng/μL nandrolone (Tokyo Chemical Industry, Japan), which was an internal standard. After centrifugation, the organic solvent was collected and dried at 40°C under the nitrogen stream and then dissolved in acetonitrile/Milli-Q water (50:50). Reversed-phase high-performance liquid chromatography (HPLC) was used to measure the amount of metabolites, as previously described.^12,13,23,24^ HPLC was performed using an Alliance 2695 separation module (Waters, Milford, MA) and a Cosmosil 5C18-MS-II Packed Column (4.6 x 150 mm, Nacalai Tesque) with continuous monitoring for ultraviolet absorbance at 254 nm. The mobile phase was maintained with acetonitrile: Milli-Q water (0:100) for the first 5 min, changed to acetonitrile: Milli-Q water (50:50) with a linear increase for the next 5 min, held for 10 min, and then linearly increased to 100% acetonitrile for the next 10 min. The typical elution times for nandrolone, 17α-hydroxyprogesterone, 20α-hydroxyprogesterone, and progesterone were 14.1, 16.7, 19.6, and 24.2 min, respectively. For quantitative analyses of progesterone and its metabolites, Empower 3 software (Waters) was used to measure the integrated peak areas based on the standard curves for nandrolone, 17α-hydroxyprogesterone, 20α-hydroxyprogesterone, and progesterone. The total protein content of the homogenate was determined using the method of Bradford (Bio-Rad).

##### Analysis of ovulated oocytes

Ovulated oocytes were obtained from the oviductal ampullae on both sides of the mouse body 16 h after 5IU PMSG/hCG injection. Oocytes were incubated in mHTF medium (Kyudo, Tosu, Japan) containing 0.1 mg/ml hyaluronidase enzyme (Sigma-Aldrich) for 5 min at 37℃ to remove cumulus cells. Tiling images of each well were captured using a 10x phase contrast objective lens of an all-in-one microscope BZ-X800 (Keyence). The number and maturation of oocytes were observed in all images. Oocytes without the 1st polar body at the germinal vesicle or in metaphase I were considered immature, and those with the 1st polar body at metaphase II were considered mature, as previously described.^25^ The average diameter of an oocyte including a zona pellucida was calculated from the value of the major axis and the minor axis using an automatic-measuring application with BZ-X800. Inaccurate measurements of the average diameter were cut off using a range setting of 75 μm < average diameter < 135 μm.

##### Statistics and reproducibility

Statistical analyses were performed using GraphPad Prism version 9. Student’s two-tailed t tests were used for statistical comparisons of data obtained under the 2 conditions. One-way analysis of variance (ANOVA) followed by Tukey’s multiple comparison test was used for the statistical comparison of more than 2 groups. All data are expressed as the means ± SE. Statistical significance was set at p < 0.05. Sample sizes and numbers are indicated in detail in the legends of each figure.

##### Data and code availability

- RNA-seq and SuperSAGE data generated in this study have been deposited at GEO under accession number GSE238259 and GSE238260, and are publicly available as of the date of publication.

Any additional information required to reanalyze the data reported in this paper is available from the lead contact upon request.

