## Supplemental Figures for "*Akr1b7* functions as a master regulator in ovarian aging"

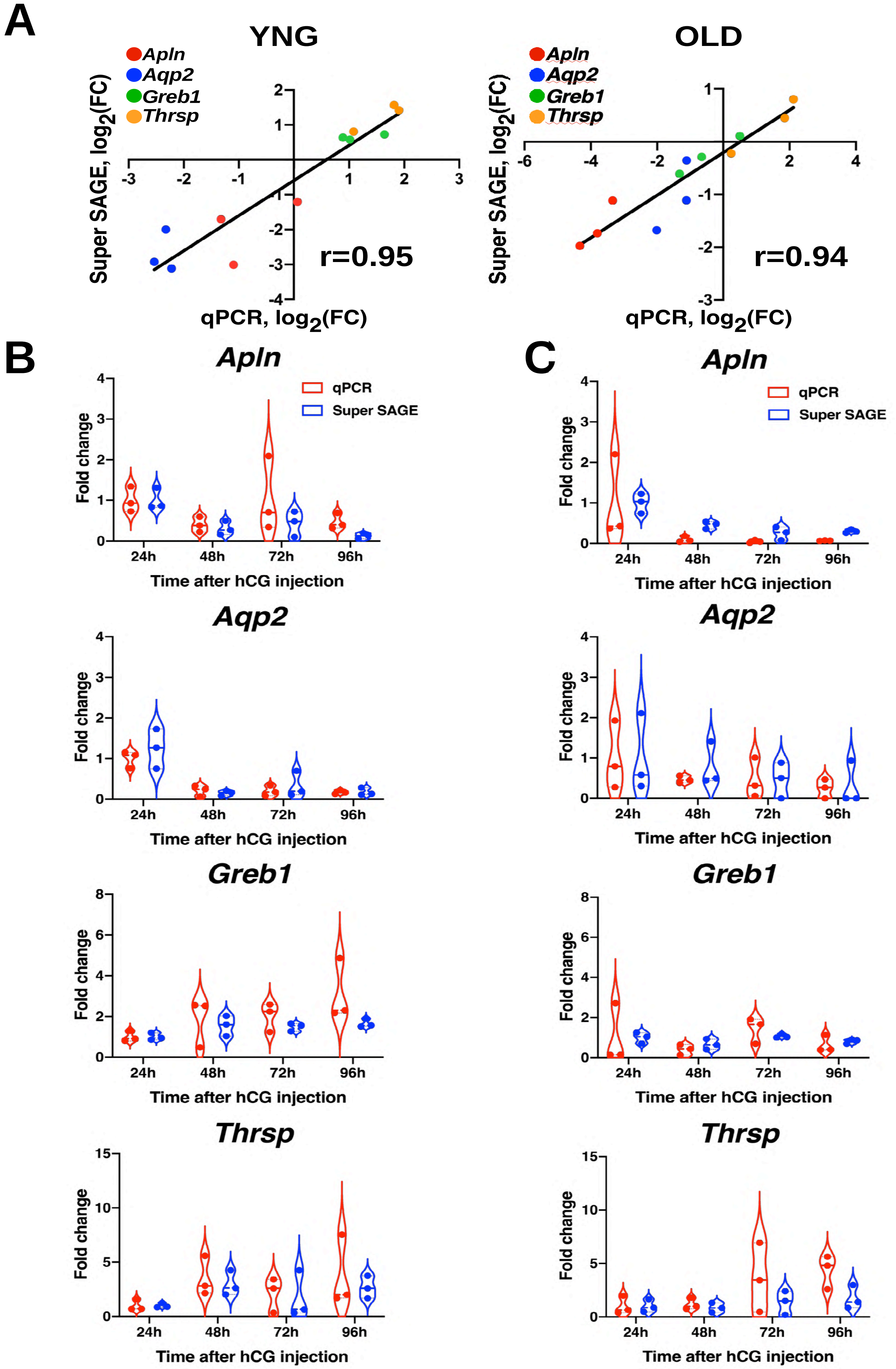
A
YNG
OLD
Apln
Aqp2
Greb1
Thrsp
Super SAGE, log2(FC)
qPCR, log2(FC)
r=0.95
r=0.94
B
C
Apln
Aqp2
Greb1
Thrsp
qPCR
Super SAGE
Fold change
24h
48h
72h
96h
Time after hCG injection

Figure S1

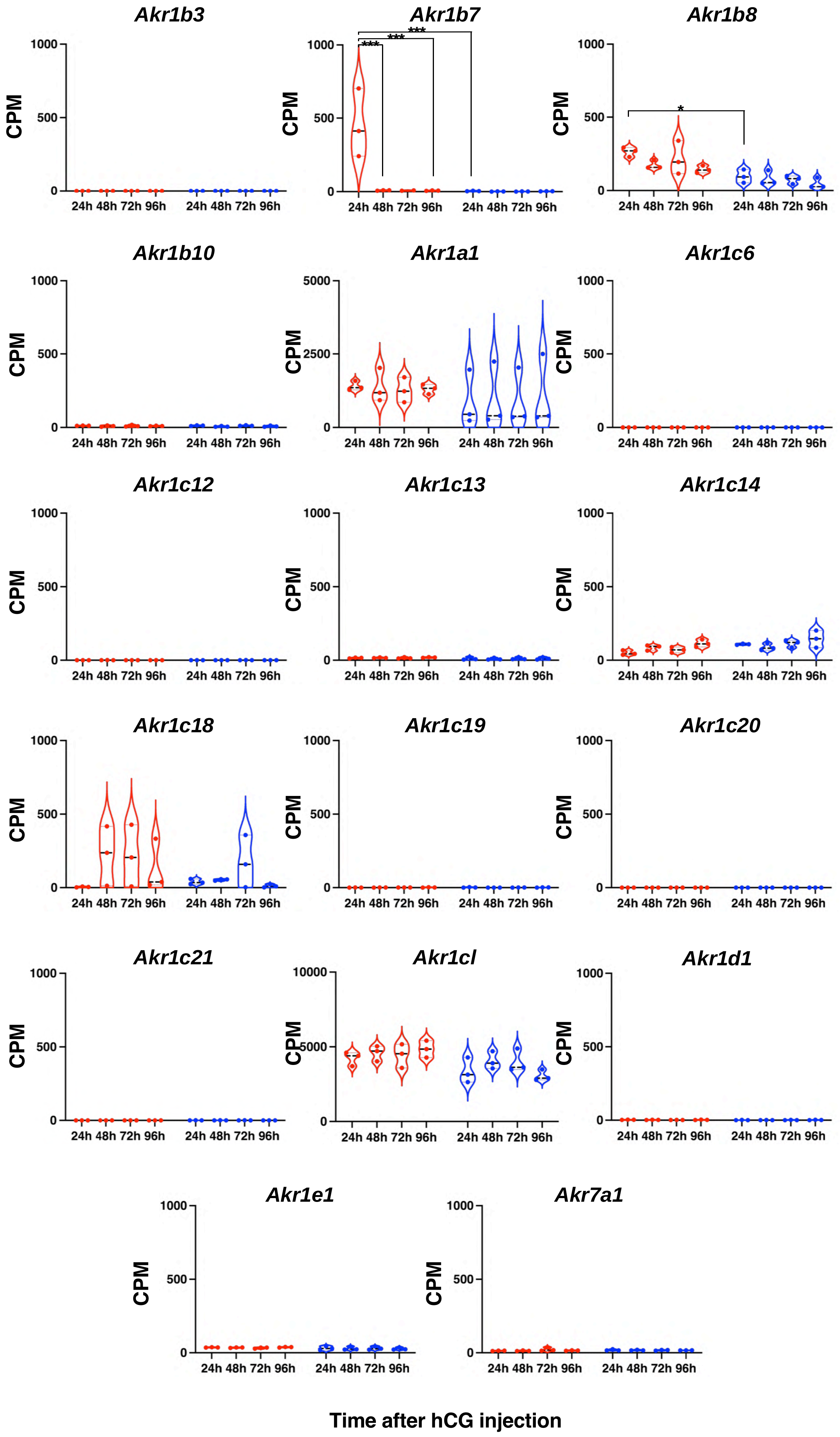

Akr1b3
Akr1b7
Akr1b8
Akr1b10
Akr1a1
Akr1c6
Akr1c12
Akr1c13
Akr1c14
Akr1c18
Akr1c19
Akr1c20
Akr1c21
Akr1cl
Akr1d1
Akr1e1
Akr7a1
CPM
24h 48h 72h 96h
Time after hCG injection


### Figure S2

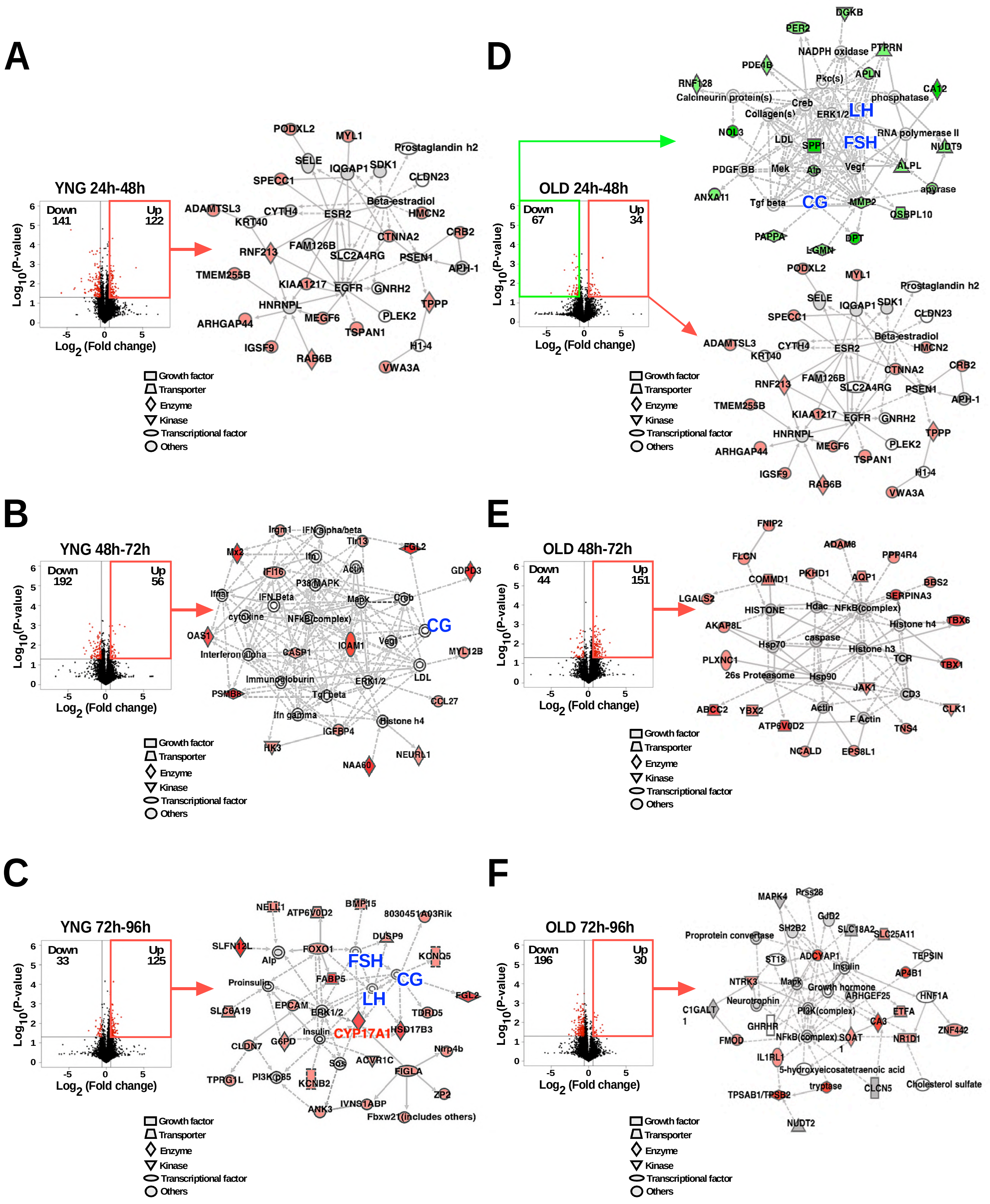
A
YNG 24h-48h
Down 141
Up 122
Log10(P-value)
Log2 (Fold change)
PODXL2
MYL1
Prostaglandin h2
SELE
IQGAP1
SDK1
CLDN23
SPECC1
Beta-estradiol
ADAMTSL3
CYTH4
ESR2
HMCN2
KRT40
CTNNA2
CRB2
RNF213
FAM126B
SLC2A4RG
PSEN1
APH-1
TMEM255B
KIAA1217
EGFR
GNRH2
HNRNPL
TPPP
ARHGAP44
MEGF6
PLEK2
TSPAN1
IGSF9
H1-4
RAB6B
VWA3A
Growth factor
Transporter
Enzyme
Kinase
Transcriptional factor
Others
D
OLD 24h-48h
Down 67
Up 34
DGKB
PER2
NADPH oxidase
PTPRN
PDE4B
Pkc(s)
APLN
RNF128
CA12
Calcineurin protein(s)
Creb
phosphatase
Collagen(s)
ERK1/2
LH
NOL3
LDL
SPP1
FSH
RNA polymerase II
NUDT9
PDGF BB
Mek
Alp
Vegf
ALPL
apyrase
ANXA11
Tgf beta
CG
MMP2
OSBPL10
PAPPA
DPT
LGMN
B
YNG 48h-72h
Down 192
Up 56
Irgm1
IFN alpha/beta
Tlr13
Mx2
FGL2
Ifn
IFI16
Actin
GDPD3
P38 MAPK
Ifnar
IFN Beta
Mapk
Creb
cytokine
NFkB(complex)
CG
OAS1
ICAM1
Vegf
Interferon alpha
CASP1
MYL12B
LDL
Immunoglobulin
ERK1/2
PSMB8
Tgf beta
CCL27
Ifn gamma
Histone h4
IGFBP4
HK3
NEURL1
NAA60
E
OLD 48h-72h
Down 44
Up 151
FNIP2
ADAM8
FLCN
PPP4R4
COMMD1
PKHD1
AQP1
BBS2
LGALS2
SERPINA3
HISTONE
Hdac
NFkB(complex)
AKAP8L
Histone h4
TBX6
caspase
Hsp70
Histone h3
PLXNC1
TCR
TBX1
26s Proteasome
Hsp90
JAK1
CD3
ABCC2
YBX2
Actin
CLK1
ATP6V0D2
F Actin
TNS4
NCALD
EPS8L1
C
YNG 72h-96h
Down 33
Up 125
NELL1
ATP6V0D2
BMP15
8030451A03Rik
DUSP9
SLFN12L
Alp
FOXO1
KCNQ5
FSH
Proinsulin
FABP5
CG
FGL2
SLC6A19
EPCAM
LH
ERK1/2
TDRD5
Insulin
CYP17A1
HSD17B3
CLDN7
G6PD
Nlrp4b
Sos
ACVR1C
TPRG1L
PI3K p85
KCNB2
FIGLA
ZP2
ANK3
IVNS1ABP
Fbxw21 (includes others)
F
OLD 72h-96h
Down 196
Up 30
MAPK4
Prss28
GJB2
Proprotein convertase
SH2B2
SLC18A2
SLC25A11
ST18
ADCYAP1
TEPSIN
Insulin
AP4B1
NTRK3
Mapk
Growth hormone
HNF1A
C1GALT1
Neurotrophin
ARHGEF25
PI3K(complex)
ETFA
GHRHR
CA3
ZNF442
FMOD
NFkB(complex)
SOAT1
NR1D1
IL1RL1
5-hydroxyeicosatetraenoic acid
TPSAB1/TPSB2
tryptase
CLCN5
Cholesterol sulfate
NUDT2

### Figure S3

**A**

| Target Sequence | PAM | Score | #Mismatch | Gene | Locus |
|---|---|---|---|---|---|
| **TTTGGTACTGAGTTCCACGA** | **AGG** | **N/A** | | **AKR1B7** | **chr6:-34412419** |
| TATG-TAGTGAGTTCCACGA | CAG | 3 | 3 | | chr7:-27554148 |
| ACTGGTAGTGAGTTCCATGA | GGG | 41 | 4 | | chr19:-23673265 |
| TTTGGTATTTAGTTCCACAA | AAG | 45 | 3 | | chr1:-79820094 |
| TTTCATAGTGAGTTCCATGA | TAG | 45 | 4 | | chr10:+82918708 |
| TTTGGTAGTGAGTTCCAGGA | AAG | 49 | 2 | | chr3:+143583509 |
| TTTCCTAGTGAGTTCCAAGA | CAG | 49 | 4 | | chr11:+86435373 |
| TATGTTAGTGAGTTCCAGGA | AAG | 49 | 4 | | chr18:+25159255 |
| CATGGTATTGAGTTCCAGGA | TAG | 49 | 4 | | chr4:-136036662 |
| CTTTGTAGTGAGTTCCAGGA | CAG | 49 | 4 | | chr7:+73625083 |
| TCTAGTAGTGAGTTCCAGGA | CAG | 49 | 4 | | chr7:-79583749 |

**B**

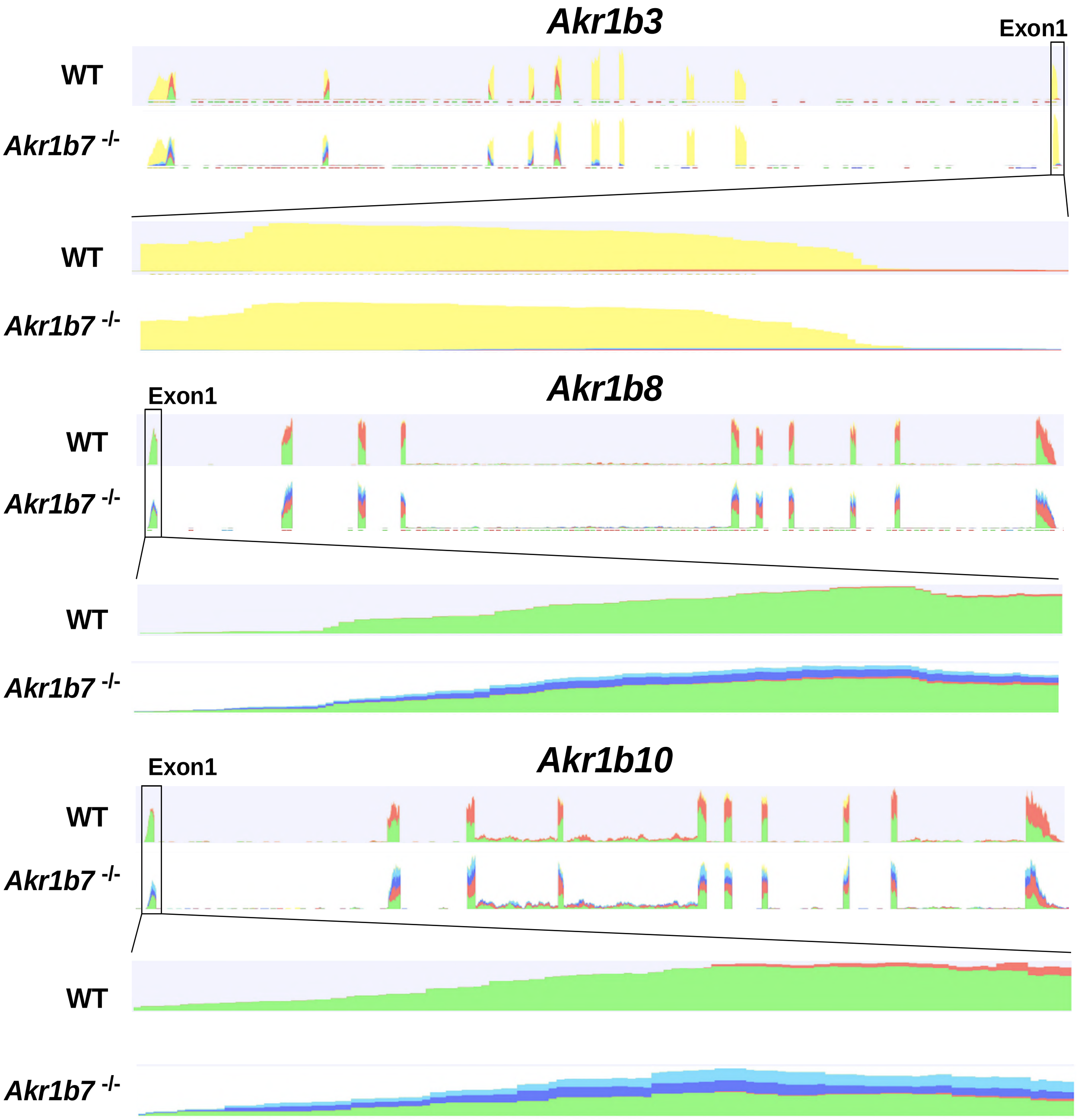


### Figure S4

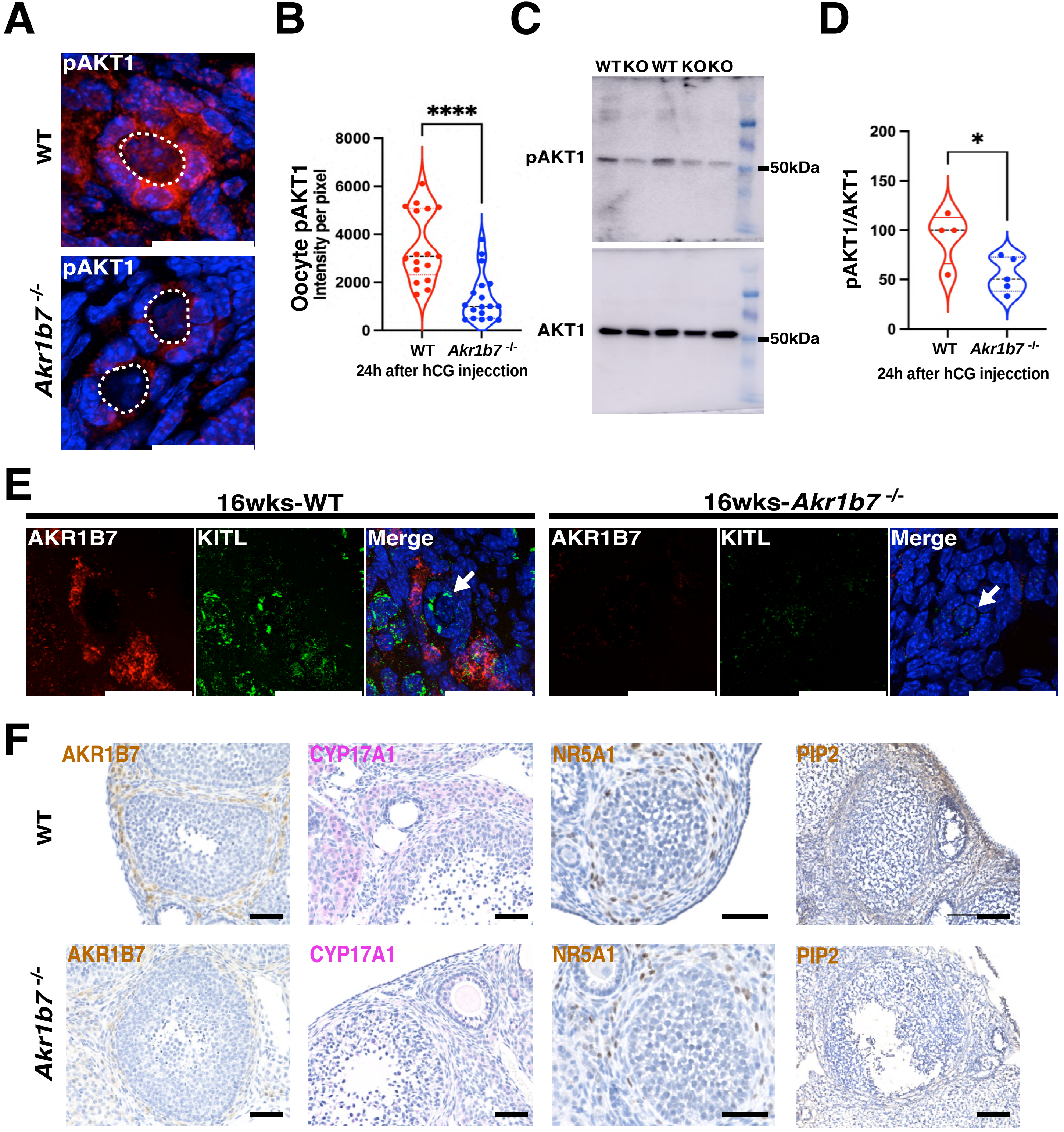
A
pAKT1
WT
pAKT1
Akr1b7 -/-
B
Oocyte pAKT1
Intensity per pixel
****
8000
6000
4000
2000
0
WT
Akr1b7 -/-
24h after hCG injecction
C
WT KO WT KO KO
pAKT1
50kDa
AKT1
50kDa
D
pAKT1/AKT1
*
200
150
100
50
0
WT
Akr1b7 -/-
24h after hCG injecction
E
16wks-WT
16wks-Akr1b7 -/-
AKR1B7
KITL
Merge
AKR1B7
KITL
Merge
F
WT
AKR1B7
CYP17A1
NR5A1
PIP2
Akr1b7 -/-
AKR1B7
CYP17A1
NR5A1
PIP2

### Figure S5

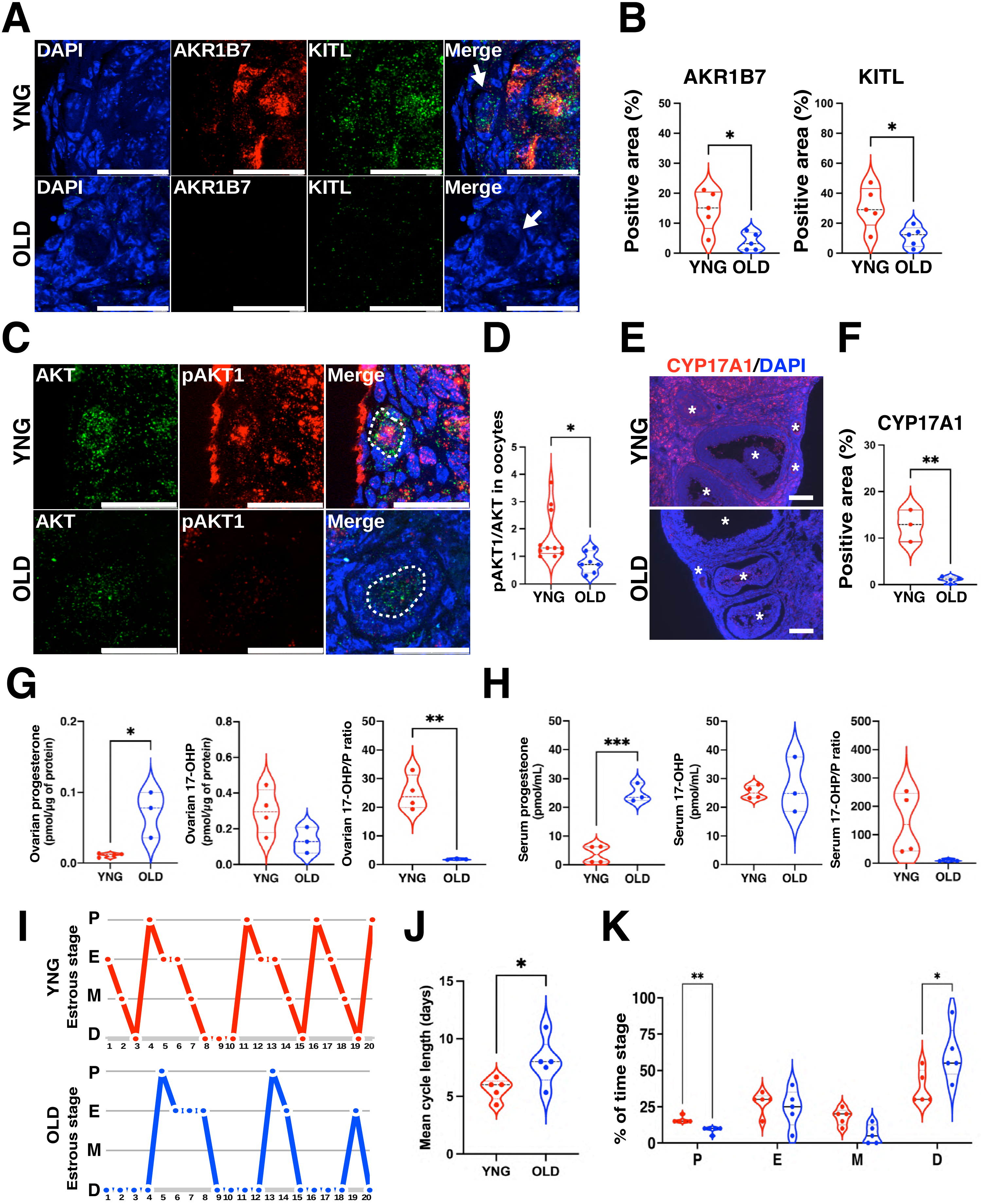

A
DAPI
AKR1B7
KITL
Merge
YNG
OLD
B
AKR1B7
KITL
Positive area (%)
YNG OLD
C
AKT
pAKT1
Merge
D
pAKT1/AKT in oocytes
E
CYP17A1/DAPI
F
CYP17A1
G
Ovarian progesterone (pmol/µg of protein)
Ovarian 17-OHP (pmol/µg of protein)
Ovarian 17-OHP/P ratio
H
Serum progesterone (pmol/mL)
Serum 17-OHP (pmol/mL)
Serum 17-OHP/P ratio
I
Estrous stage
J
Mean cycle length (days)
K
% of time stage
P
E
M
D


Figure S6

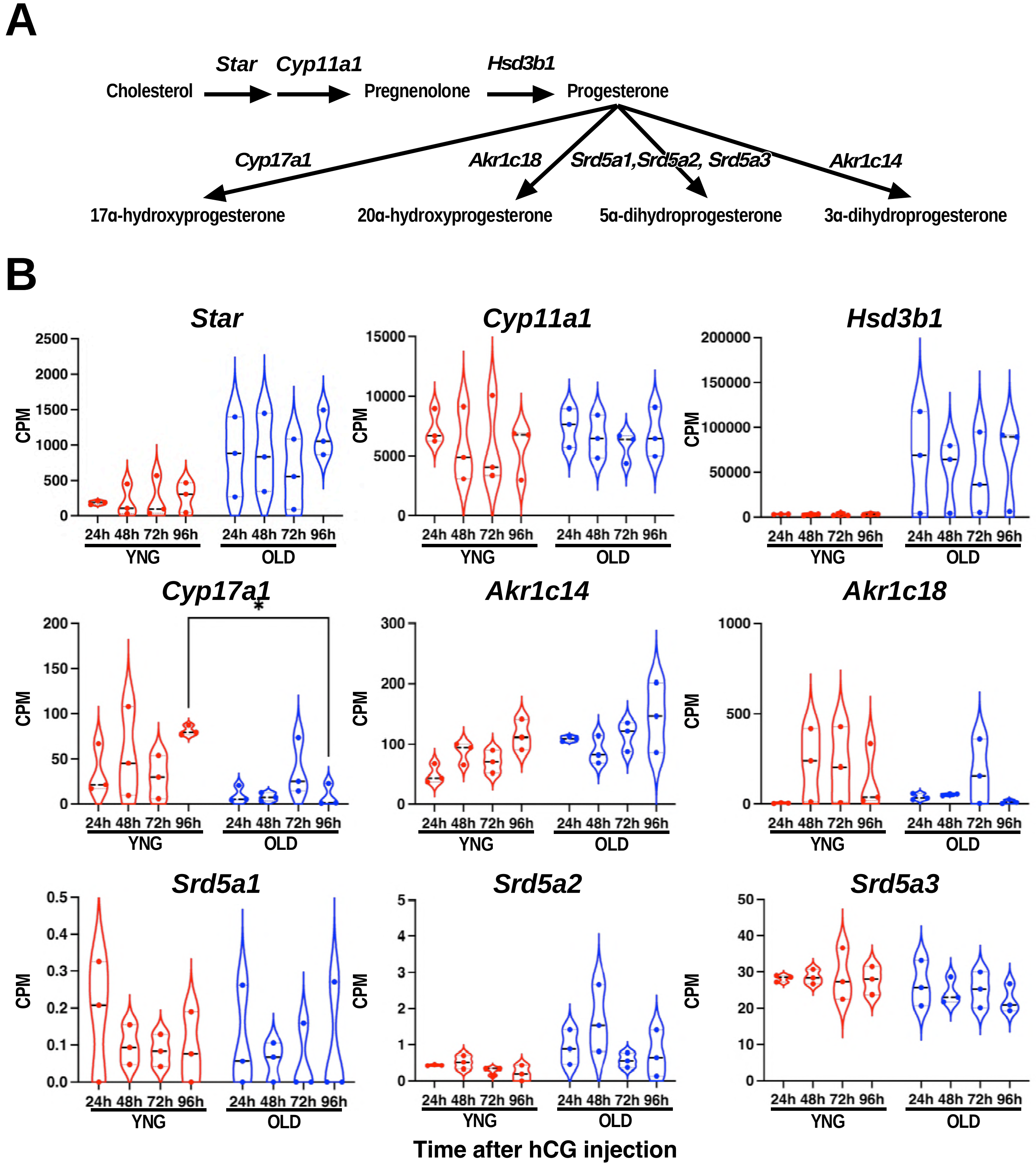
A
Star
Cyp11a1
Hsd3b1
Cholesterol
Pregnenolone
Progesterone
Cyp17a1
Akr1c18
Srd5a1,Srd5a2, Srd5a3
Akr1c14
17α-hydroxyprogesterone
20α-hydroxyprogesterone
5α-dihydroprogesterone
3α-dihydroprogesterone
B
Star
Cyp11a1
Hsd3b1
Cyp17a1
Akr1c14
Akr1c18
Srd5a1
Srd5a2
Srd5a3
CPM
24h 48h 72h 96h
YNG
OLD
Time after hCG injection

Figure S7
